# Chitin perception in plasmodesmata identifies subcellular, context-specific immune signalling in plants

**DOI:** 10.1101/611582

**Authors:** Cecilia Cheval, Matthew Johnston, Sebastian Samwald, Xiaokun Liu, Annalisa Bellandi, Andrew Breakspear, Yasuhiro Kadota, Cyril Zipfel, Christine Faulkner

## Abstract

The plasma membrane (PM) that lines plasmodesmata has a distinct protein and lipid composition, underpinning specific regulation of these connections between cells. The plasmodesmal PM can integrate extracellular signals differently from the cellular PM, but it is not known how this specificity is established or how a single stimulus can trigger independent signalling cascades in neighbouring membrane domains. Here we have used the fungal elicitor chitin to investigate signal integration and responses at the plasmodesmal PM. We found that the plasmodesmal PM employs a receptor complex composed of the LysM receptors LYM2 and LYK4 which respectively change their location and interactions in response to chitin. Downstream, signalling is transmitted via a specific phosphorylation signature of an NADPH oxidase and localised callose synthesis that causes plasmodesmata closure. This demonstrates the plasmodesmal PM deploys both plasmodesmata-specific components and differential activation of PM-common components to independently integrate an immune signal.

## Introduction

Plants have an interconnected cytoplasm that bridges neighbouring cells throughout. Plasma membrane-lined tubes called plasmodesmata (singular plasmodesma) cross the cell wall and establish cytoplasmic, plasma membrane (PM) and endoplasmic reticulum (ER) continuity between cells. At a single cell interface there might be thousands of plasmodesmata (Faulkner *et al.*, 2008; Danila *et al.*, 2018) and as small molecules, proteins and RNAs can pass from cell-to-cell through plasmodesmata this establishes considerable capacity for direct molecular exchange between cells and tissues. As almost all cells within plants are connected via plasmodesmata, they create an extensive conduit through which signals and resources can be distributed between cells, tissues and organs.

The functional aperture of a plasmodesma can change in response to a range of developmental and environmental signals, generating dynamic zones of connectivity and molecular exchange that underpin whole organism responses (Rinne *et al.*, 2011; Faulkner *et al.*, 2013; Lim *et al.*, 2016; Rinne, Paul and van der Schoot, 2018). Changes to plasmodesmal aperture can be executed by the synthesis and hydrolysis of the β-1,3-glucan callose in the cell wall that surrounds a plasmodesma (Rinne *et al.*, 2011; Vaten *et al.*, 2011; Cui and Lee, 2016). Enzymes responsible for callose turnover are anchored in the plasmodesmal PM and their activity is likely regulated by actors that similarly reside in the plasmodesmal PM. Indeed, several plasmodesmata-located proteins have been shown to be critical for plasmodesmal responses to a range of signals (Benitez-Alfonso *et al.*, 2013; Faulkner *et al.*, 2013; Wang *et al.*, 2013; Cui and Lee, 2016). Further, there are examples of proteins, such as the Arabidopsis CRINKLY 4 (ACR4) and CLAVATA 1 (CLV1) receptor kinases (RKs), that are resident in both the PM and the plasmodesmal PM but form plasmodesmata-specific complexes (Stahl *et al.*, 2013) suggesting that the plasmodesmal PM is a unique signalling environment.

The PM is highly organised into microdomains that are diverse in composition, function and dynamics. These microdomains regulate the organisation and activation of a variety of proteins resident within the plasma membrane. In plants and animals, this includes a variety of cell-surface proteins involved in the perception of environmental and developmental signals (Malinsky *et al.*, 2013; Ott, 2017; Gronnier *et al.*, 2018). One such class of cell surface proteins are receptors: RKs and receptor proteins (RPs) display receptor domains on the outward facing side of the PM and are critical for detecting changes in the extracellular environment and initiating downstream signalling. Plants have a large repertoire of these PM-anchored receptors and can recognise a large range of molecular signals. Some RKs and RPs can dynamically form modular receptor complexes in PM microdomains and/or nanodomains, forming signalling hubs that execute response outputs (Lingwood and Simons, 2010; Bucherl *et al.*, 2017; Ott, 2017). For example, Medicago LysM-CONTAINING RECEPTOR-LIKE KINASE 3 (LYK3) is dynamically recruited to a PM microdomain during rhizobia infection (Liang *et al.*, 2018) and the flg22 receptor FLAGELLIN SENSING 2 (FLS2) is stabilised in nanodomains during signalling (Bucherl *et al.*, 2017). The protein and lipid composition of the plasmodesmal PM differs from other domains of the PM (Fernandez-Calvino *et al.*, 2011; Grison *et al.*, 2015; Tilsner *et al.*, 2016); the plasmodesmal PM is rich in sphingolipids (Grison *et al.*, 2015) suggesting it has similarity to PM microdomains in its lipid composition, further supporting the hypothesis that the plasmodesmal PM hosts specific signalling cascades similar to PM microdomains.

Here we have elucidated the mechanism of plasmodesmal PM specific signalling events and determined that the plasmodesmal PM functions independently of the neighbouring PM. By examining the mechanism by which cells perceive and respond to chitin, a fungal cell wall component that triggers immune responses, we have determined that signalling in the plasmodesmal PM is distinct from signalling in the PM. We found that in addition to the plasmodesmata located receptor LysM-CONTAINING GPI-ANCHORED PROTEIN 2 (LYM2) (Faulkner *et al.*, 2013), chitin responses in the plasmodesmal PM require two additional LysM RKs, LYK4 and LYK5. However, only LYK4 is detected in plasmodesmata. LYM2 can associate with LYK4 suggesting that chitin-triggered plasmodesmal responses are mediated by a chitin-activated LYM2-LYK4 complex. The dependence of plasmodesmal responses on LYK5 appears to rest on LYK5-dependent modification of LYK4 in the PM prior to chitin perception. Chitin perception by LYM2-LYK4 triggers BOTRYTIS INDUCED KINASE 1 (BIK1)-independent, calcium-dependent protein kinase mediated phosphorylation of the NADPH oxidase RESPIRATORY BURST OXIDASE HOMOLOG PROTEIN D (RBOHD) and ultimately induces callose deposition and plasmodesmata closure. These findings identify that the plasmodesmal PM integrates extracellular signals independently and specifically relative to the PM, supporting a model in which the plasmodesmal PM hosts discrete signalling cascades. Further, the independence of chitin-triggered plasmodesmal PM responses from those in the PM suggest that cell-to-cell connectivity is regulated independently of other immune responses. This work demonstrates that plasmodesmata are regulated independently of other immune responses, suggesting that there is a critical requirement for a cell to finely tune connectivity to its neighbours.

## Results

### Chitin-triggered plasmodesmata closure is dependent on LYK4 and LYK5

We previously identified that LYM2 is a GPI-anchored, LysM receptor protein that is resident in the plasmodesmal PM (Faulkner *et al.*, 2013). As LYM2 has no intracellular domains we reasoned that it must interact with other proteins to initiate downstream signals that result in plasmodesmal responses. Ligand perception by LysM RKs and RPs often involves multiple members of the LysM protein family: chitin perception in rice involves both the RP CHITIN ELICITOR BINDING PROTEIN (OsCEBiP) and the RK CHITIN ELICITOR RECEPTOR KINASE 1 (OsCERK1) (Kaku *et al.*, 2006; Hayafune *et al.*, 2014); peptidoglycan perception in Arabidopsis involves the RK CERK1, and RPs LYM1 and LYM3 (Willmann *et al.*, 2011); and PM chitin perception in Arabidopsis involves CERK1 (also called LYK1) and the RKs LYK4 and LYK5 (Cao *et al.*, 2014). Thus, we hypothesised that LYM2 might partner with a LysM RK for signalling. The Arabidopsis LysM RK family consists of 5 members: CERK1/LYK1, LYK2, LYK3, LYK4 and LYK5. To narrow down plasmodesmata signalling candidates we screened publicly available data sets for LYK gene expression. Comparing data sets from seedlings (GSE74955, Yamada *et al*., 2016; GSE78735, Hillmer *et al*., 2017) and mature leaves (eFP browser, Winter *et al*. 2007) we identified variable expression patterns for the *LYK* family members (Fig. S1). Thus, we performed RT-PCR to identify members of the family expressed in mature Arabidopsis leaves where we assay for and detect LYM2 function. Only transcripts from *CERK1*, *LYK3*, *LYK4* and *LYK5* were detected in mature leaves grown in our conditions, eliminating LYK2 from further analysis (Fig. S1). We previously showed that CERK1 is not required for chitin-triggered plasmodesmata closure (Faulkner *et al.*, 2013) and therefore we assayed for chitin-triggered plasmodesmal responses in *lyk3*, *lyk4* and *lyk5-2* mutants. Microprojectile bombardment assays, in which movement of GFP from single transformed cells within the Arabidopsis leaf epidermis is measured, demonstrated that *lyk3* mutants show a chitin-triggered reduction in the spread of GFP, indicative of plasmodesmata closure (Fig. 1) comparable to the wild-type (Col-0) response. By contrast, *lyk4* and *lyk5-2* mutants show no change in GFP spread following chitin treatment identifying that these mutants cannot close their plasmodesmata in response to chitin similar to *lym2* mutant plants. Thus, LYK4 and LYK5 are required for chitin-triggered plasmodesmata closure.

**Figure 1.**
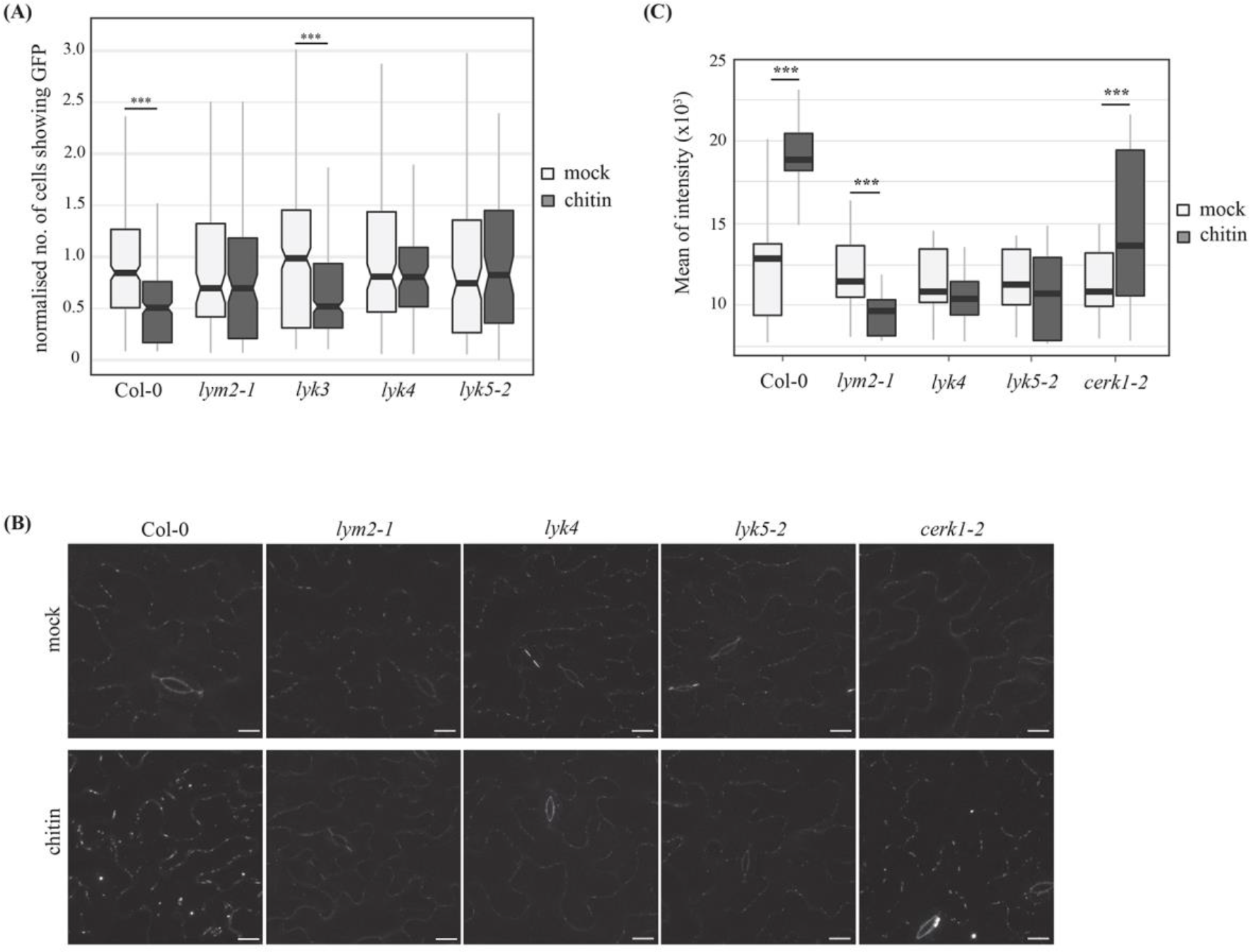
LYK4 and LYK5 regulate plasmodesmal permeability in response to chitin. (A) Microprojectile bombardment into leaf tissue of 5–6‐week‐old Arabidopsis shows that Col-0 and *lyk3* but not *lym2-1*, *lyk4* and *lyk5-2* exhibit reduced flux of GFP to neighbouring cells in response to chitin. Data was collected from 6 biological replicates and the number of cells showing GFP has been normalised to the mean of the mock-treated data within genotypes. This data is summarised in box-plots in which the line within the box marks the median, the box signifies the upper and lower quartiles, the whiskers represent the minimum and maximum within 1.5 × interquartile range. Notches represent approximate 95% confidence intervals. Asterisks indicate statistical significance compared to control conditions (n ≥ 85, ***p-value < 0.001). (B) Confocal images of aniline blue stained plasmodesmal callose in leaves of 5–6‐week‐old Col-0 plants, as well as *lym2-1, lyk4, lyk5-2,* and *cerk1-2* mutants. Images were acquired 30 min post-infiltration with water or chitin. Scale bars are 15 μm. (C) Quantification of plasmodesmata-associated fluorescence of aniline blue stained callose using automated image analysis. Col-0 and the *cerk1-2* mutant show an increase in aniline blue stained plasmodesmal callose 30 min post-chitin treatment. In *lym2-1, lyk4* and *lyk5-2* this response is not detected. This correlates with the flux phenotype and identifies that chitin-triggered plasmdesmata closure is caused by callose deposition at plasmodesmata. The fluorescence intensity is summarised in box-plots in which the line within the box marks the median, the box signifies the upper and lower quartiles, the minimum and maximum within 1.5 × interquartile range. Data was analysed by pairwise Students’ t-test comparing mock to chitin treated tissue within genotypes (n ≥ 31, ***p-value < 0.001).

### LYM2-dependent chitin-signalling induces callose deposition at plasmodesmata

Callose is a β-1,3-glucan polymer deposited at plasmodesmata during stages of development and in response to a range of stresses inducing functional plamodesmal closure (Wang *et al.*, 2013; Cui and Lee, 2016; Xu *et al.*, 2017). It is established that reactive oxygen species (ROS) induce plasmodesmal callose deposition and recently we showed that callose is deposited at plasmodesmata in response to flg22 (Xu *et al.*, 2017). Therefore, we examined callose deposition at plasmodesmata in response to chitin in Arabidopsis to determine if this is common to pathogen-triggered plasmodesmata closure. We quantified aniline blue-stained plasmodesmata-located callose deposits in chitin-treated and mock-treated leaf tissue of WT plants and, as for flg22, plasmodesmata-located aniline blue fluorescence increased significantly within one hour of chitin treatment (Fig. 1B, C). This indicates that callose deposition at plasmodesmata in response to chitin is an early immune response, likely triggered within the timeframe of other rapid PAMP-triggered responses such as the ROS burst and MAPK activation. Aniline blue staining of *lym2-1*, *lyk4* and *lyk5-2* mutant leaves in the presence of chitin showed that each mutant is unable to deposit callose at plasmodesmata in response to chitin, confirming that the chitin-triggered reduction in GFP movement through plamsodesmata is caused by callose deposition. *lym2-1* mutant leaves showed a reduction in plasmodesmal callose in response to chitin. Like Col-0, *cerk1-2* mutants show an increase in plasmodesmal callose deposition in response to chitin, consistent with the chitin-triggered reduction in GFP movement through plasmodesmata by the bombardment assay.

### LYK4 is present in plasmodesmata

Having identified that LYK4 and LYK5 are required for chitin-triggered plasmodesmata closure we examined their subcellular localisation to determine if they accumulate at plasmodesmata like LYM2. We generated translational fusions of LYK4 and LYK5 to RFP. Both LYK4 and LYK5 localised to the PM in the absence and presence of chitin, with no enrichment in plasmodesmata evident (Fig. S2). Plasmodesmata are smaller than the limits of resolution of a confocal microscope and if there is no enrichment of protein at the plasmodesmata relative to the PM it is difficult to conclude absolutely whether the protein is present or absent from plasmodesmata. Thus, we performed subcellular fractionation of plasmodesmata and protein extraction to determine the presence or absence of LYK4 and LYK5 in plasmodesmal fractions (Fig. 2A). For this we expressed LYK4-HA, LYK5-HA or the plasmodesmal protein PDLP5-HA (Lee *et al.*, 2011) in *N. benthamiana* tissue and purified plasmodesmata. We could detect PDLP5-HA and LYK4-HA, in purified plasmodesmal extracts, but not LYK5-HA, identifying that only LYK4 co-locates with LYM2 at plasmodesmata. This suggests that despite the genetic dependence of chitin-triggered plasmodesmata closure on *LYM2*, *LYK4* and *LYK5*, these three receptors do not necessarily act co-operatively within plasmodesmata.

**Figure 2.**
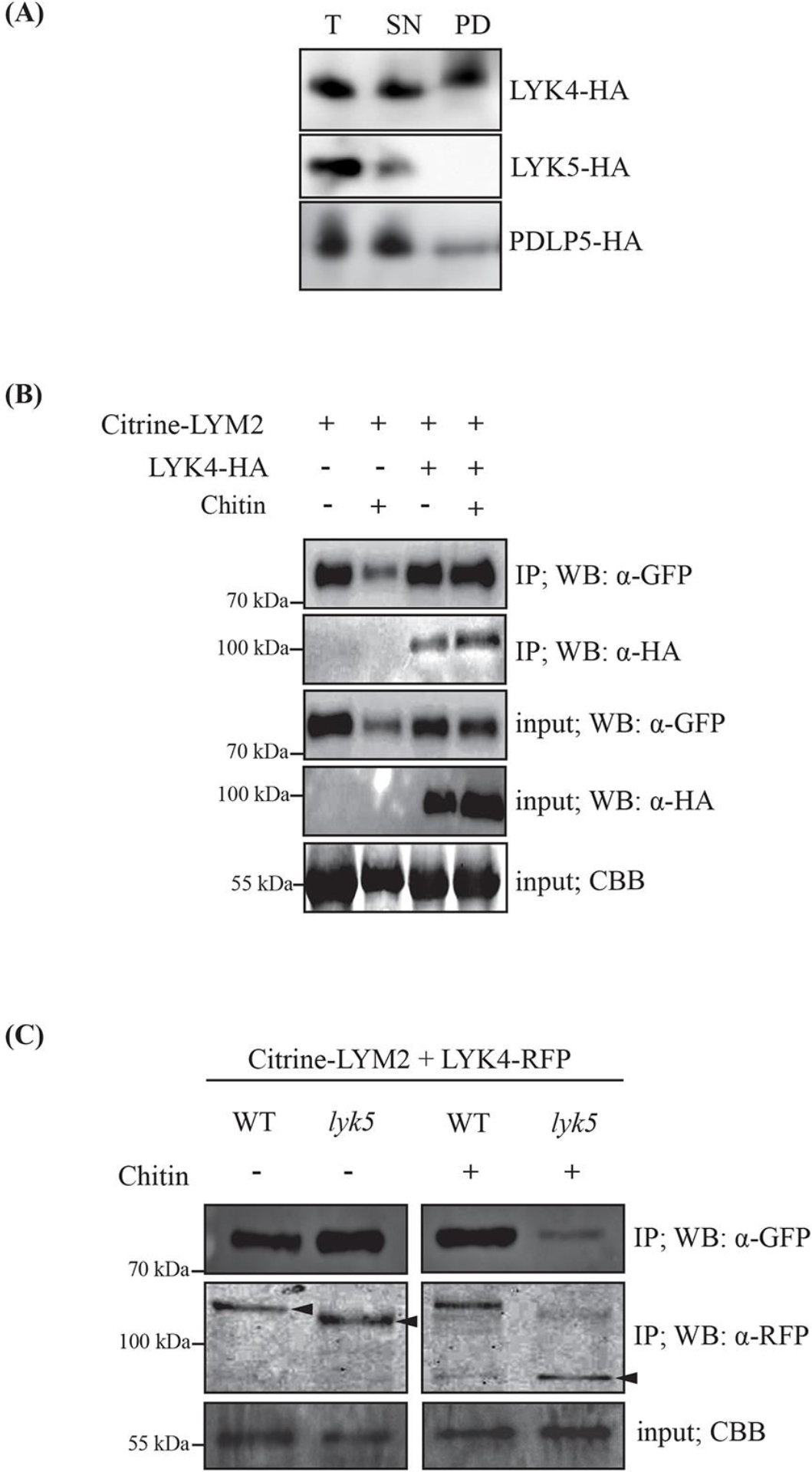
LYK4 is detected in plasmodesmata and associates with LYM2. (A) Western blot analysis of purified plasmodesmata fractions from *N. benthamiana* tissue co-expressing LYK4-HA, LYK5-HA or PDLP5-HA. Total (T; extracts from ground tissue), supernatant (SN; all cellular material excluding cell walls) and plasmodesmatal (PD; membranes released from purified cell walls) extracts were separated by SDS-PAGE and Western blots were probed with anti-HA and to determine the presence of LYK4-HA, LYK5-HA and PDLP5-HA in each fraction. Experiments were repeated four times with similar results. (B) Western blot analysis of immunoprecipitated protein extracts from *N. benthamiana* tissue expressing Citrine-LYM2 and LYK4-HA. LYK4-HA is detected in samples purified from membrane fractions by immunoprecipitation of Citrine-LYM2 with GFP-trap beads. Input and immunoprecipitated (IP) samples were probed α-GFP and α-HA antibodies as indicated. CBB, Coomassie brilliant blue. Size standards are indicated to the left of the panel. Experiments were repeated three times with similar results. (C) Western blot analysis of immunoprecipitated protein extracts from Arabidopsis protoplasts expressing Citrine-LYM2 and LYK4-HA. LYK4-HA is detected in samples purified from membrane fractions by immunoprecipitation of Citrine-LYM2 with GFP-trap beads from both Col-0 and *lyk5-2* protoplasts. Input and immunoprecipitated (IP) samples were probed α-GFP and α-HA antibodies as indicated. LYK4-HA bands of different sizes are indicated by arrowheads. CBB, Coomassie brilliant blue. Size standards are indicated to the left of the panel. Experiments were repeated three times with similar results.

### LYM2 associates with LYK4

The presence of LYM2 and LYK4 in the plasmodesmal PM suggests that they may form a functional complex that triggers plasmodesmal responses. To test this hypothesis, we determined if LYM2 associates with LYK4 by targeted co-immunoprecipitation (co-IP) assays. We co-expressed Citrine-LYM2 with LYK4-HA in *N. benthamiana* and immunoprecipitated Citrine-LYM2 from membrane fractions with anti-GFP beads (Fig. 2B). LYK4-HA was co-immunoprecipitated with LYM2 from both water and chitin treated tissue suggesting that LYM2 associates with LYK4 in a chitin independent manner. The negative control BRI1-RFP did not co-immunoprecipitate with Citrine-LYM2 (Fig. S3A).

Given that LYM2 and LYK4 are present in plasmodesmata we thought that these two proteins could directly execute plasmodesmal PM chitin signalling. However, LYK5 is also required for chitin-triggered plasmodesmata closure and thus, we tested the dependence of the interaction between LYM2 and LYK4 on LYK5. For this we transformed Arabidopsis *lyk5-2* protoplasts with Citrine-LYM2 and LYK4-RFP (Fig. 2C) and performed targeted co-IPs. We observed that LYK4-RFP immunoprecipitated with Citrine-LYM2 in a chitin-independent manner from both Col-0 and *lyk5-2* protoplasts. While this clearly demonstrates that the interaction between LYM2 and LYK4 is LYK5 independent, these assays also identified that LYK4-RFP appears approximately 10 kDa smaller on SDS-PAGE gels when extracted from *lyk5-2* protoplasts, suggesting that LYK4 is modified in a LYK5 dependent manner. Further, following chitin treatment the dominant LYK4-RFP band detected in *lyk5-2* protoplasts appears approximately 30kDa smaller than the dominant band in the WT protoplasts suggesting that the ‘non-modified’ variant of LYK4-RFP in the *lyk5-2* mutant is subject to degradative processing in response to chitin. Given the genetic dependence of chitin-triggered plasmodesmata closure on *LYK5*, these observations suggest that modification of LYK4 mediated by LYK5 is essential for its plasmodesmata-related signalling function.

### LYK4 and LYK5 associate in the PM

The observation that LYK5 is required for both chitin-triggered plasmodesmata closure and modification of LYK4, but is not located at plasmodesmata, suggests a model in which LYK5 plays a spatio-temporally distinct role in chitin-triggered plasmodesmata closure. To further investigate we explored the possibility that LYK5 associates with and modifies LYK4 in the PM, upstream of LYK4 activity at plasmodesmata. Pull-downs of LYK4-GFP from membrane fractions of *N. benthamiana* tissue expressing both LYK4-GFP and LYK5-RFP showed that LYK5-RFP co-immunoprecipitates with LYK4-GFP (Fig. 3A). The negative control BRI1-RFP did not co-immunoprecipitate with LYK4-GFP (Fig. S3B). This suggests that LYK4 and LYK5 associate in the PM.

**Table 1.**
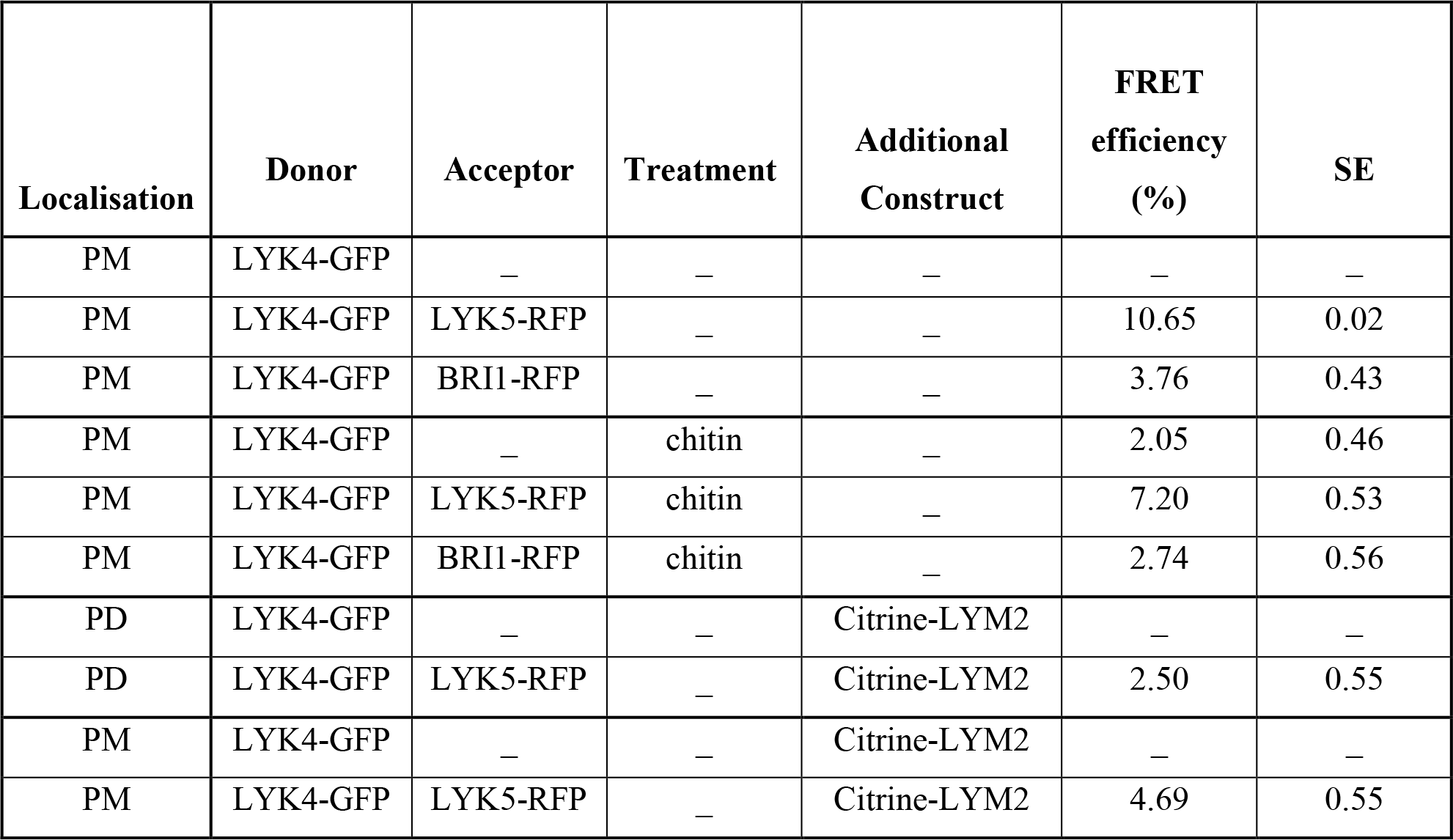
FRET efficiencies of LYK4-GFP in the presence and absence of the acceptor-RFP. The mean of FRET efficiency is expressed in percentage (%). PM, plasma membrane; PD, plasmodesmata; SE, standard error.

| Localisation | Donor | Acceptor | Treatment | Additional Construct | FRET efficiency (%) | SE |
| --- | --- | --- | --- | --- | --- | --- |
| PM | LYK4-GFP | – | – | – | – | – |
| PM | LYK4-GFP | LYK5-RFP | – | – | 10.65 | 0.02 |
| PM | LYK4-GFP | BRI1-RFP | – | – | 3.76 | 0.43 |
| PM | LYK4-GFP | – | chitin | – | 2.05 | 0.46 |
| PM | LYK4-GFP | LYK5-RFP | chitin | – | 7.20 | 0.53 |
| PM | LYK4-GFP | BRI1-RFP | chitin | – | 2.74 | 0.56 |
| PD | LYK4-GFP | – | – | Citrine-LYM2 | – | – |
| PD | LYK4-GFP | LYK5-RFP | – | Citrine-LYM2 | 2.50 | 0.55 |
| PM | LYK4-GFP | – | – | Citrine-LYM2 | – | – |
| PM | LYK4-GFP | LYK5-RFP | – | Citrine-LYM2 | 4.69 | 0.55 |

**Table 2.**
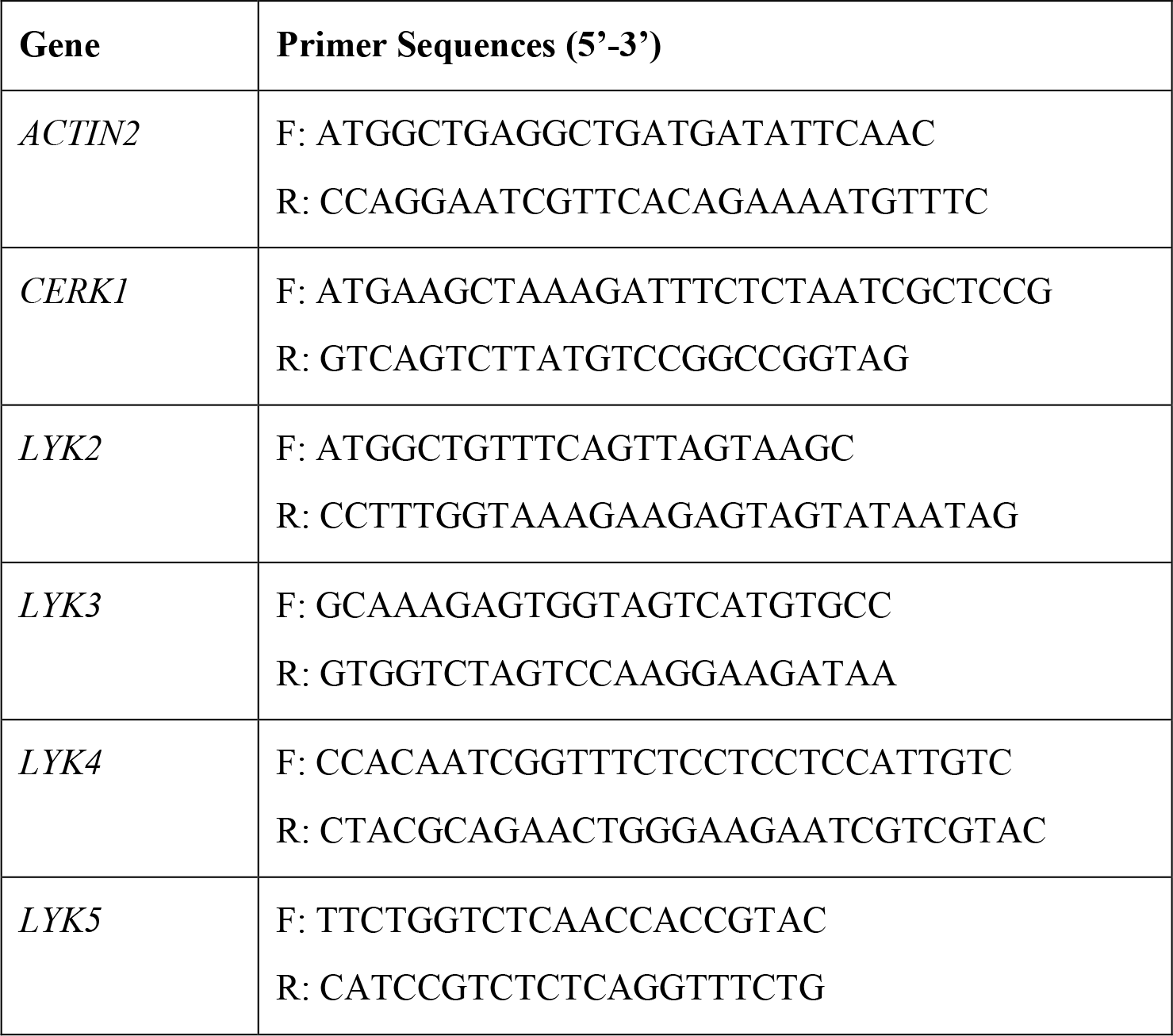
Primer sequences used in RT-PCR analysis of *LYSM-RK*s.

**Figure 3.**
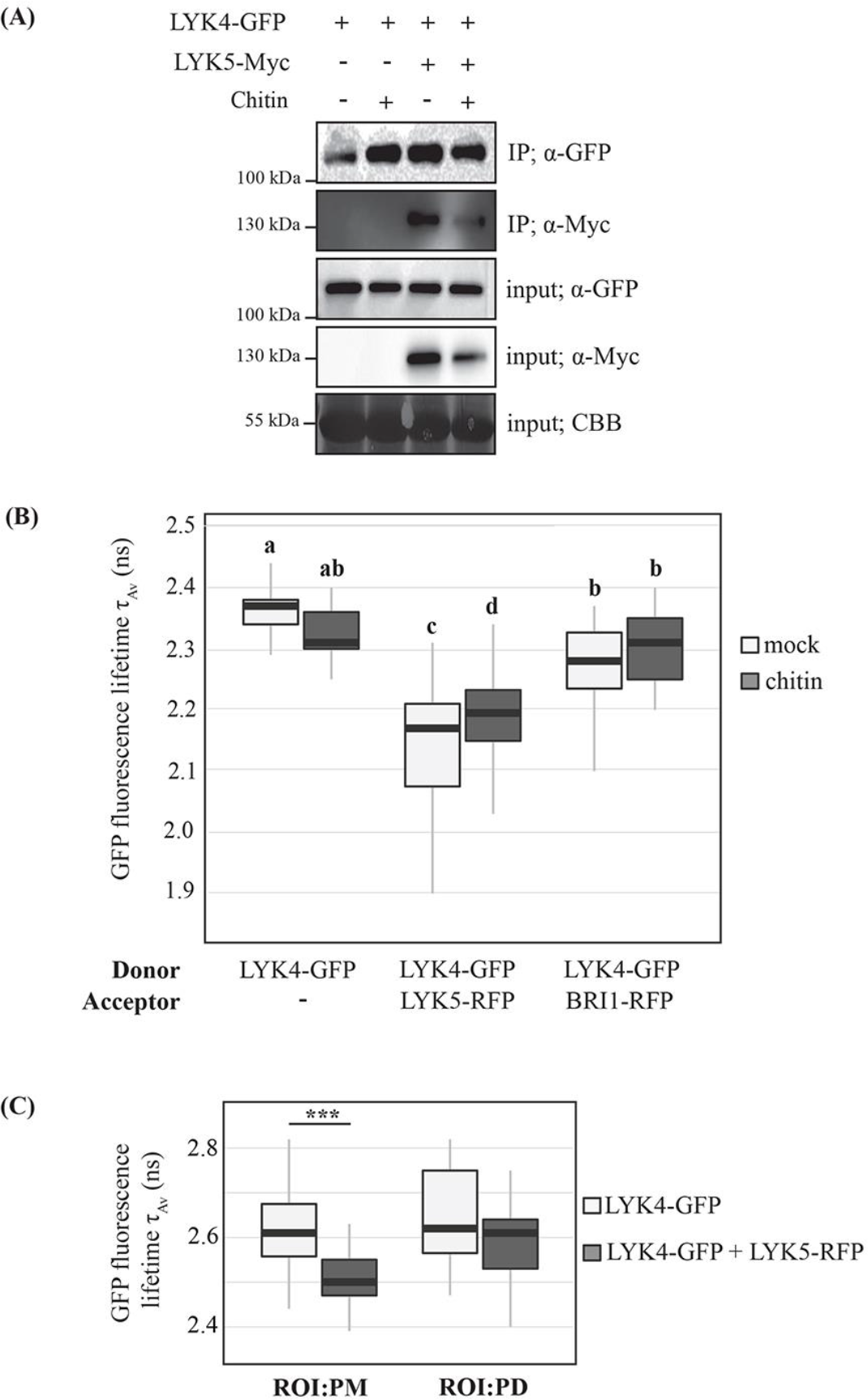
LYK5 dynamically associates with LYK4 in the PM. (A) Western blot analysis of immunoprecipitated protein extracts from *N. benthamiana* tissue expressing LYK4-GFP and LYK5-Myc. LYK4-GFP was immunoprecipitated from membrane fractions and western blots of input and immunoprecipitated extracts were probed with α-GFP and α-Myc to detect LYK4-GFP and LYK5-Myc respectively. Coomassie brilliant blue (CBB) stained membranes serve as a loading control. Size standards are indicated to the left of the panel. Experiments were repeated three times with similar results. (B) FRET-FLIM analysis of LYK4-GFP in the presence of acceptors LYK5-RFP or BRI1-RFP, and the presence and absence of chitin. Fluorescence lifetime was measured in *N. benthamiana* tissue transiently co-expressing the indicated constructs as donors or acceptors. Box-plots represent GFP fluorescence-weighted average lifetime (τ_Av_, ns) fitted with a double-exponential model: the line within the box marks the median, the box signifies the upper and lower quartiles, the whiskers represent the minimum and maximum within 1.5 × interquartile range. Data was analysed by ANOVA with a post-hoc Tukey multiple comparison of means (p-value <0.01, n ≥19). Samples with the same letter code are not significantly different. (C) FRET-FLIM analysis of LYK4-GFP at the plasmodesmal PM and the PM in the presence and absence of LYK5-RFP RFP. Fluorescence lifetime was measured in in *N.benthamiana* tissue transiently co-expressing the noted constructs. Plasmodesmata were marked by co-expression of Citrine-LYM2 and ROI were defined around plasmodesmata (PD) and in the PM for analysis. Box-plots represent GFP fluorescence-weighted average lifetime (τ_Av_, ns) fitted with a double-exponential model: the line within the box marks the median, the box signifies the upper and lower quartiles, the whiskers represent the minimum and maximum within 1.5 × interquartile range. Data was analysed by a Students’ t-test in which the lifetime of LYK4-GFP was compared in the absence and presence of LYK5-RFP for both ROIs. Asterisks indicate statistical significance compared to control conditions (n≥ 27, p-value < 0.001).

We further explored the dynamics of this interaction with FLIM-FRET analysis (Fig. 3B). The fluorescence lifetime (average amplitude, Τ_Av_) of PM-localised LYK4-GFP was significantly reduced in the presence of LYK5-RFP, indicating an increase in FRET as expected for interacting proteins. Chitin-treatment decreased the degree of FRET observed between LYK4-GFP and LYK5-RFP suggesting that chitin weakened the interaction between LYK4 and LYK5 by either complex dissociation or a change in conformation.

We also used FRET-FLIM to investigate the localisation of a LYK4-LYK5 complex and compare the PM with the plasmodesmal PM (Fig. 3C). For this we marked plasmodesmata by co-expression of LYK4-GFP and LYK5-RFP with Citrine-LYM2. When we compared Τ_Av_ of LYK4-GFP in regions of interest (ROIs) within the PM and ROIs at plasmodesmata we observed that in the PM Τ_Av_ was significantly reduced by the presence of LYK5-RFP as expected, but that in plasmodesmata it was not reduced. This supports our finding that LYK5 is not present in the plasmodesmal PM, and the hypothesis that LYK5 is critical for LYK4 function upstream of plasmodesmal signalling.

### LYM2, LYK4 and LYK5 are dynamic in response to chitin

It has been shown that LYK5 undergoes re-localisation in the form of endocytosis in response chitin (Erwig *et al*., 2017) and so we explored possible dynamics of LYM2 in response to chitin. Live-cell imaging of Citrine-LYM2 in *N. benthamiana* leaves suggested that Citrine-LYM2 fluorescence at plasmodesmata increased in response to chitin treatment (Fig. 4A); i.e. Citrine-LYM2 at plasmodesmata appeared to get brighter relative to PM fluorescence following chitin treatment. We quantified this response by measuring the plasmodesmal index (PD index) of fluorescence (plasmodesmata:PM fluorescence intensity (Perraki *et al.*, 2018) for LYM2 in the absence and presence of chitin (Fig. 4B). The mean PD index of Citrine-LYM2 in mock treated samples was 1.7 but following 30 min chitin treatment increased to 2.5, indicating a significant increase in fluorescence intensity at plasmodesmata in response to chitin. Fluorescence anisotropy measurements of Citrine-LYM2 in chitin-treated tissue identified lower anisotropy in the plasmodesmata relative to the PM indicating there is more homo-FRET occurring at plasmodesmata (Fig 4C). The observation that anisotropy (*r*) of Citrine-LYM2 is lower in the PM relative to freely rotating cytosolic GFP suggests that homo-FRET occurs in the PM, *i.e.* that LYM2 can interact with itself in the PM. That *r* is further reduced in plasmodesmata suggests that Citrine-LYM2 forms a higher order complex or signalling platform in the plasmodesmal PM.

**Figure 4.**
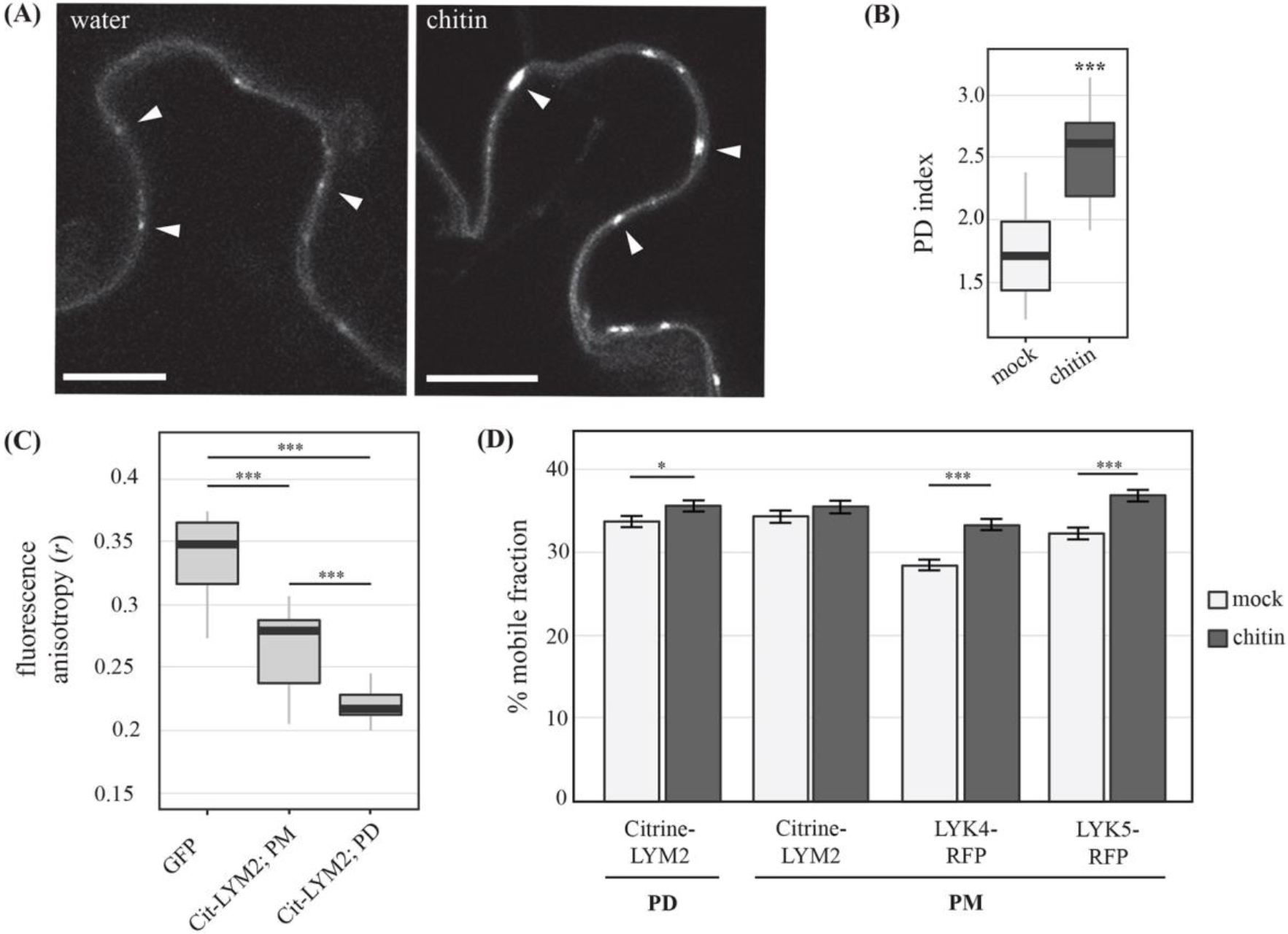
LYM2 accumulates at PD in response to chitin. (A) Single plane confocal images of *N.benthamiana* tissue expressing Citrine-LYM2 before and after chitin treatment. Left panel shows Citrine-LYM2 in water-treated tissue and the right shows Citrine-LYM 30 min post chitin treatment. Arrowheads indicate example plasmodesmata. Scale bars, 10 μm. (B) Quantification of the PD index of Citrine-LYM2 at PD in *N.benthamiana* after mock and chitin treatments. Data was analysed with a Students t-test (***p-value < 0.001, n=31). (C) Fluorescence anisotropy (*r*) of cytosolic GFP, PM located Citrine-LYM2 and plasmodesmata-located (PD) Citrine-LYM2. Data was analysed pairwise by a Students’ t-test (***p-value <0.001, GFP: n=11, Citrine-LYM2: n=21). (D) % mobile fraction of LYM2, LYK4 and LYK5 as measured by FRAP assays. For Citrine-LYM2 FRAP measurements were taken for the plasmodesmata-located (PD) and PM-located pools of receptor. For LYK4-RFP and LYK5-RFP FRAP measurements were taken in the PM. Data was analysed by LOESS regression and estimated marginal means analysis, error bars are standard error (***p-value <0.001, n ≥43).

We did not observe any robust change in the distribution of fluorescence of either LYK4-RFP or LYK5-RFP in response to chitin by live-imaging in *N. benthamiana* leaves (Fig. S2). However, FRET-FLIM data suggests that LYK4 and LYK5 dissociate in response to chitin, which could lead to increased lateral mobility of LYK4 and LYK5. To test this, we performed FRAP analysis of LYK4-RFP and LYK5-RFP in mock and chitin-treated tissue (Fig. 4D). Following chitin treatment, the mobile fraction of both LYK4 and LYK5 within the PM increased. Citrine-LYM2 also exhibited a marginal increase in the mobile fraction of molecules in response to chitin, but only at plasmodesmata. This elevation in the proportion of the mobile fraction of each receptor in response to chitin suggest dynamic behaviour of these receptors that changes their interactions and/or the membrane domains in which they reside.

### LYM2-dependent chitin-triggered plasmodesmata closure engages unique calcium/ROS regulatory modules

Reactive oxygen species (ROS) are produced in immune responses and can induce plasmodesmata closure (Torres, Dangl and Jones, 2002; Cui and Lee, 2016). Thus, we hypothesized that ROS play a role in the regulation of plasmodesmata closure in response to chitin, downstream of LYM2-LYK4 activity. We firstly verified that H_2_O_2_ -induced plasmodesmata closure can be detected by the bombardment method and observed a reduction of GFP movement from cell-to-cell in WT plants after H_2_O_2_ treatment (Fig. S4). In this assay, *lym2-1* mutants were able to close their plasmdesmata in response to H_2_O_2_ demonstrating that any role for ROS signalling in chitin-triggered plasmodesmata closure occurs independently, or downstream of LYM2 activity. We also further established that transient expression of LYM2, LYK4 and LYK5 enhanced ROS production in *N. benthamiana* leaf discs following chitin treatment, suggesting that each of these LysM proteins can mediate ROS signalling.

The rapid production of ROS in response to chitin is associated with the NADPH oxidase RBOHD (Kadota *et al.*, 2014) and so we tested whether RBOHD is required for chitin-triggered plasmodesmata closure. Bombardment assays showed that the *rbohd* mutant is not able to close its plasmodesmata in response to chitin treatment (Fig. 5A). This was supported by quantitative analysis of plasmodesmal callose that showed that *rbohd* mutants did not deposit callose at plasmodesmata in response to chitin (Fig. S5). Thus, ROS produced by RBOHD are a critical component of the chitin-triggered signalling cascade that induces plasmodesmata closure.

**Figure 5.**
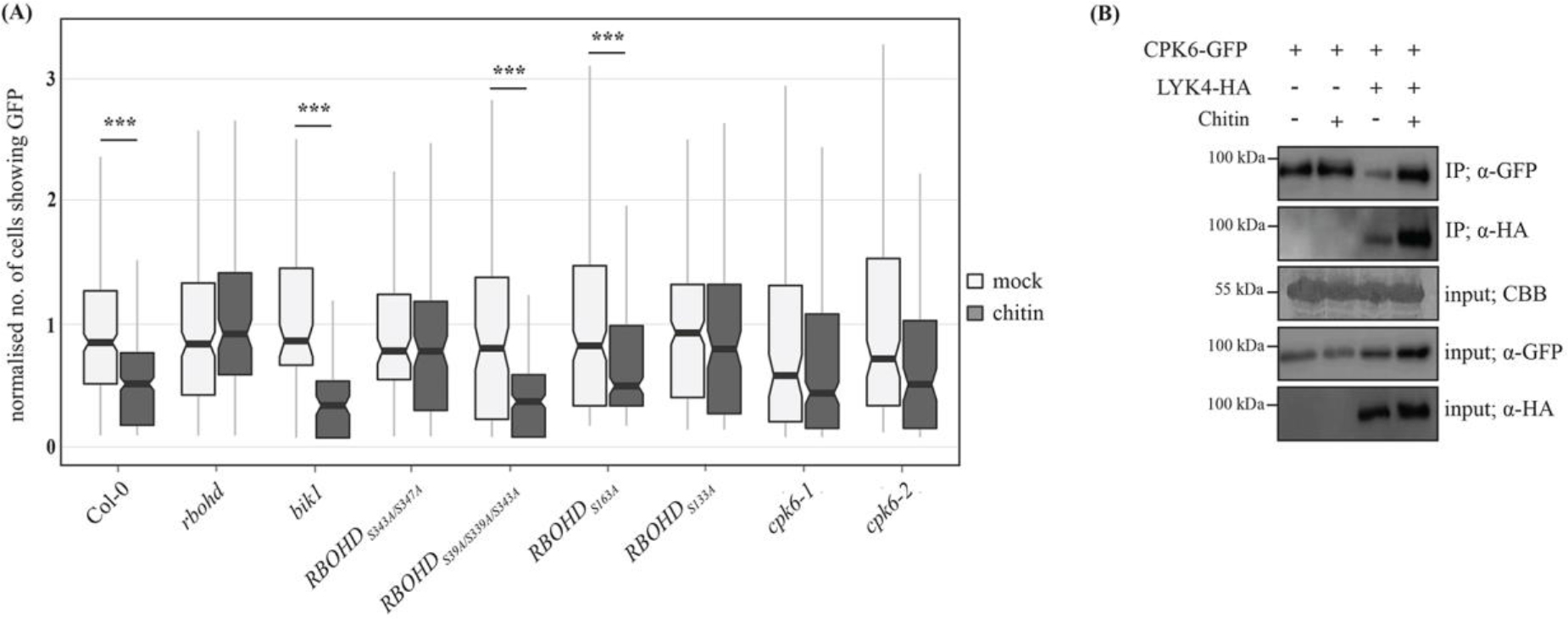
CPK-dependent phosphorylation of RBOHD is required for plasmodesmata closure in response to chitin. (A) Microprojectile bombardment into leaf tissue shows that *rbohd* mutants do not show a reduction in GFP flux to neighbouring cells in response to chitin. The *bik1* mutant, and RBOHD phosphosite mutant variants *S39A/S339A/S343A* and *S163A* exhibit a similar reduction in movement of GFP to neighbouring cells in response to chitin. By contrast, the RBOHD phosphosite mutant variants *S343A/S347A* and *S133A*, and the *CPK6* mutants *cpk6-1* and *cpk6-2* show no reduction in GFP flux to neighbouring cells in response to chitin. Data was collected from 6 biological replicates and the number of cells showing GFP has been normalised to the mean of the mock data within genotypes. This data is summarised in box-plots in which the line within the box marks the median, the box signifies the upper and lower quartiles, the minimum and maximum within 1.5 × interquartile range. Notches represent approximate 95% confidence intervals. Data has been analysed by a Students’ t-test, or by a Mann-Whitney U-test for non-normal data (n ≥ 93, ***p-value < 0.001). (B) LYK4 co-immunoprecipitates with CPK6 in *N. benthamiana*. CPK6-GFP was transiently co-expressed with LYK4-HA in *N. benthamiana* leaf tissue and CPK6-GFP was immunoprecipitated using GFP-trap beads. Input and immunoprecipitated (IP) samples were analysed by western blots probed with α-GFP and α-HA. Coomassie brilliant blue (CBB) staining of membranes serves as a loading control. The experiment was repeated three times with similar results.

In Arabidopsis, RBOHD is regulated by phosphorylation by protein kinases (Dubiella *et al.*, 2013; Kadota *et al.*, 2014, 2018; Li *et al.*, 2014). It has been shown that the receptor-like cytoplasmic kinase (RLCK) BIK1 and calcium dependent protein kinases (CPKs) both phosphorylate RBOHD in response to chitin: BIK1 phosphorylates Ser39, Ser339, Ser343 and Ser347, while CPKs phosphorylate Ser133, Ser148, Ser163 and Ser347 (Dubiella *et al.*, 2013; Kadota *et al.*, 2014). To test dependence of plasmodesmal responses on BIK1, we assayed *bik1* mutant plants for chitin-triggered plasmodesmata closure and observed that *bik1* mutants exhibited a wild-type response to chitin. Thus, BIK1 does not function in this process (Fig. 5A). Given the large number of members in the RLCK and CPK families that function in immune responses we focused our attention on the contribution of phosphorylated residues of RBOHD to chitin-triggered plasmodesmal responses. For this we tested a range of phospho-null mutant variants (mutation of serine to alanine) of RBOHD in the *rbohd* mutant background: RBOHD_S39A/S339A/S343A_, RBOHD_S343A/S347A_, RBOHD_S133A_, and RBOHD_S163A_ (Fig. 5A). Consistent with our results for the *bik1* mutant, RBOHD_S39A/S339A/S343A_, in which serines phosphorylated by BIK1 are mutated, showed a wild-type response and can close their plasmodesmata in response to chitin. Examining variants in which mutated phosphosites are targeted by CPKs, RBOHD_S343A/S347A_ and RBOHD_S133A_ were unable to close their plasmodesmata in response to chitin while RBOHD_S163A_ showed a wild-type response, identifying that phosphorylation of S347 and S133 are required for chitin-triggered plasmodesmata closure. Thus, unlike the CERK1-mediated ROS burst at the PM (Miya *et al.*, 2007; Rao *et al.*, 2018), chitin-triggered plasmodesmata closure is independent of BIK1 and instead dependent on CPK mediated phosphorylation of RBOHD.

Among the CPKs, CPK4, 5, 6 and 11 have been described to be involved in the regulation of RBOHD-dependent ROS burst, and CPK6 has been shown to phosphorylate both Ser133 and S347 (Kadota *et al.*, 2014). Therefore, we tested the involvement of CPK6 in chitin-triggered plasmodesmata closure. Bombardment assays showed that two independent *cpk6* mutants, *cpk6-1* and *cpk6-2*, were unable to close their plasmodesmata in response to chitin (Fig. 5A) confirming CPK6 functions in plasmodesmal signalling.

LYM2 is a GPI anchored protein and therefore unlikely to directly interact with intracellular CPKs. Having shown previously that LYM2 associates with LYK4 (Fig 2), we investigated the interaction between CPK6 and LYK4. We generated a GFP-tagged translational fusion of CPK6 (CPK6-GFP) that we transiently expressed in *N. benthamiana* leaf tissue with LYK4-HA. Co-IP experiments show that CPK6-GFP and LYK4-HA associate (Fig 5B). These data suggest that a LYM2-LYK4 complex recruits CPK6 for phosphorylation of RBOHD and thereby executes plasmodesmal signalling.

## Discussion

The innate immune system depends on the perception of pathogen molecules by extracellular and intracellular receptors. Here, we have characterised how a plant cell can perceive and initiate immune responses to a single ligand in different domains of the PM. Our data has identified that while fungal chitin is perceived and triggers signalling via a CERK1-dependent cascade in the PM, in the plasmodesmal PM chitin triggers a LYM2/LYK4-dependent cascade to produce a specific, localised response. Thus, for chitin signalling in plant cells both receptors and signalling cascades can show specificity to a given subcellular context.

Subcellular specificity in ligand perception by receptors and signalling have both been documented in animal cells. It is known that the same ligand can be perceived by different receptors in different subcellular compartments, allowing detection of a ligand in different cellular locations, e.g. the flg22 peptide can be perceived extracellularly by the membrane anchored Toll-Like Receptor 5 (TLR5) (Hayashi *et al.*, 2001) and in the cytosol by the soluble NLR Family CARD Domain Containing 4 (NLRC4) (Kofoed and Vance, 2011; Zhao *et al.*, 2011). Alternatively, subcellular specificity can be conveyed at the response level, e.g. the lipopolysaccharide (LPS) receptor TLR4 triggers signalling from the PM but is also endocytosed and subsequently initiates a secondary signalling cascade from endosomes (Płóciennikowska *et al.*, 2015). Our findings here show that plant cells can combine both receptor and signalling specialisation to fully integrate the perception of a signal into an immune response. Further, specialisation and independence of signalling at the plasmodesmal PM suggests that cell-to-cell connectivity must be regulated independently of other immune outputs, raising questions regarding whether there is a critical requirement for cells to balance communication and exchange with the protective mechanism imposed by isolation.

Our investigation in to the mechanisms by which chitin signalling is executed at the plasmodesmal PM identified that the plasmodesmata-resident GPI-anchored protein LYM2 accumulates at plasmodesmata in response to chitin. LYM2 also exhibits greater homo-FRET in the plasmodesmal PM than in the PM suggesting it oligomerises or clusters to form a signalling platform. It is well established that many forms of immune signalling occur via the formation of supramolecular complexes such as signalosomes and inflammasomes that involve the oligomerisation of cytosolic NLR receptors and downstream signalling components. Membrane anchored receptors also form multi-component signalling complexes (Mueller and Nickel, 2012; Stegmann *et al.*, 2017; Ren *et al.*, 2019) and in some cases higher order clusters (Scott *et al.*, 2008; Pan *et al.*, 2019) for signalling. The CYSTEINE-RICH RECEPTOR-LIKE KINASE2 (CRK2) was recently shown to accumulate at the plasmodesmal PM in response to salt stress (Hunter *et al.*, 2019) suggesting the possibility that the plasmodesmal PM commonly executes signalling via transient recruitment and concentration of machinery.

We found that in addition to LYM2, the RK LYK4 is also critical for plasmodesmal PM chitin signalling. LYK4 associates with LYM2 and is detected in plasmodesmata but does not accumulate at plasmodesmata in response to chitin. LYK4 associates with the RK LYK5 in the PM but in response to chitin treatment we observed that the mobile fraction of LYK4 increased and the association between LYK4 and LYK5 in the PM decreased. When membrane proteins change their associations, they frequently exhibit a change in their biophysical properties and/or behaviour and therefore our data suggests that chitin triggers dissociation between these two RKs and releases a pool of LYK4 for signalling at plasmodesmata. It has been reported that the mobility of receptors both increases (Wang *et al.*, 2015) and decreases (Bucherl *et al.*, 2017) in response to ligand binding, but given that signalling complexes are larger multi-component structures, it is likely that activated receptors in complexes are less mobile. Our observations of an increased mobile fraction in response to the presence of chitin might result from the complex dynamics and changes in association that are evident for the LysM receptor family and represent the disassociated fraction of the RK pool. It might also be that LysM receptors and their signalling complexes behave differently from the LRR receptors that have been the focus of previous studies.

In addition to LYM2 and LYK4 we identified that LYK5 is a critical component of chitin-triggered plasmodesmal responses. However, LYK5 was not detected in purified plasmodesmata. This suggested to us that the dependence of chitin-triggered plasmodesmata closure on LYK5 is due to activity and association with LYK4 in the PM, upstream of plasmodesmal signalling responses. Targeted co-IP of LYM2 and LYK4 from *lyk5-2* protoplasts identified that LYK4 was smaller in the absence of LYK5 and undergoes enhanced degradative processing in response to chitin. It is known that LysM receptors are subject to ectodomain cleavage (Petutschnig *et al.*, 2014), which might explain the chitin-triggered degradation of LYK4. However, the smaller form of LYK4 detected in *lyk5-2* protoplasts is more consistent with a post-translational modification such as mono-ubiquitination or sumoylation. Indeed, LYK5 has been shown to interact with the E3 ubiquitin ligase PLANT U-BOX13 (Liao *et al.*, 2017) suggesting ubiquitination is a likely candidate modification. Thus, we propose that LYK5 is required for post-translational modification of LYK4 so that it can execute plasmodesmal responses.

LYK4 is predicted to be an inactive kinase (Wan *et al.*, 2012), suggesting that it needs further interacting partners to mediate signalling. Here, we have shown LYK4 can associate with CPK6, which is known to phosphorylate Ser133 and Ser347 of RBOHD (Kadota *et al.*, 2014) and is critical for chitin-triggered plasmodesmata closure. Thus, CPK6 could transmit a signal from LYK4 to RBOHD in the plasmodesmal PM. In this scenario, LYK4 has characteristics of a scaffold protein that modulates signalling complex formation: LYK4 bridges the apoplastic plasmodesmal-specificity defined by LYM2 with an intracellular signalling kinase. This has some similarity to the activity described recently for the malectin-like receptor kinase FERONIA (FER) (Stegmann *et al.*, 2017) and the N-myristoylated protein BRASSINOSTEROID SIGNALING KINASE 3 (BSK3) (Ren *et al.*, 2019) in the formation of receptor signalling complexes for EF-Tu and flg22, and brassinosteroids respectively.

Current models for phosphorylation and activation of RBOHD have suggested that RLCKs and CPKs are involved in a two-step regulation of RBOHD activity in which RLCKs prime RBOHD for activation by CPK phosphorylation (Kadota *et al.*, 2014). This idea allows for finely tuned control of the production of ROS. In our study, chitin-triggered plasmodesmata closure is independent of BIK1, one of the RLCK family members required for the CERK1 associated chitin-triggered ROS burst (Rao *et al.*, 2018). Chitin-triggered plasmodesmata closure is unlikely to depend on other RLCKs as the response is also independent of phosphorylation of Ser39, Ser339 and Ser343 which are targeted by the RLCK BIK1 and essential for the CERK1 associated chitin-triggered ROS burst (Kadota *et al.*, 2014). Thus, our data suggests that for chitin-triggered plasmodesmata closure, RBOHD activity might be controlled by different regulatory modules. It is unclear what information specific activation of RBOHD conveys in independent signalling pathways to a single ligand.

We observed that chitin-triggered plasmodesmata closure is dependent on both CPK-targeted phosphosites within RBOHD, and CPK6. However, not all CPK targeted phosphosites of RBOHD we tested were critical for plasmodesmal responses. Mutation of Ser163 did not abolish chitin-triggered plasmodesmata closure while mutations in Ser133 and Ser347 did, implying there is response-associated specificity within CPK-mediated phosphorylation of RBOHD. A possible explanation lies in CPK phosphorylation motifs (Kadota *et al.*, 2014): Ser133 and Ser347 are categorised as motif 1 sites ([B-B-X-B]-φ-X-X-X-X-S/T-X-B) while Ser163 is categorised as a motif 2 site (φ-X-B-X-X-S-X-X-X-φ) linking chitin-triggered plasmodesmata closure to motif 1 sites. The significance of these motifs and their relevance to plasmodesmal signalling is unknown; they may correlate to specific CPKs, or to an independent mechanism for tuning RBOHD activity.

Combining all the measurements and observations we have made in this study, we propose a model for plasmodesmal PM specific signal integration during chitin immune responses (Fig. 6). We suggest that LYK4 and LYK5 constitutively interact at the PM and that this mediates function-critical modification of LYK4. Chitin perception triggers dissociation of the LYK4-LYK5 complex, allowing for an increased interaction between LYK4 and LYM2, and LYM2 accumulation at plasmodesmata. At the plasmodesmal PM, LYM2 establishes a signalling platform that recruits LYK4, and by association CPK6, to this membrane microdomain. CPK6 phosphorylates RBOHD to produce ROS and induce localised callose synthesis and plasmodesmata closure within an hour of the initial perception of chitin.

**Figure 6.**
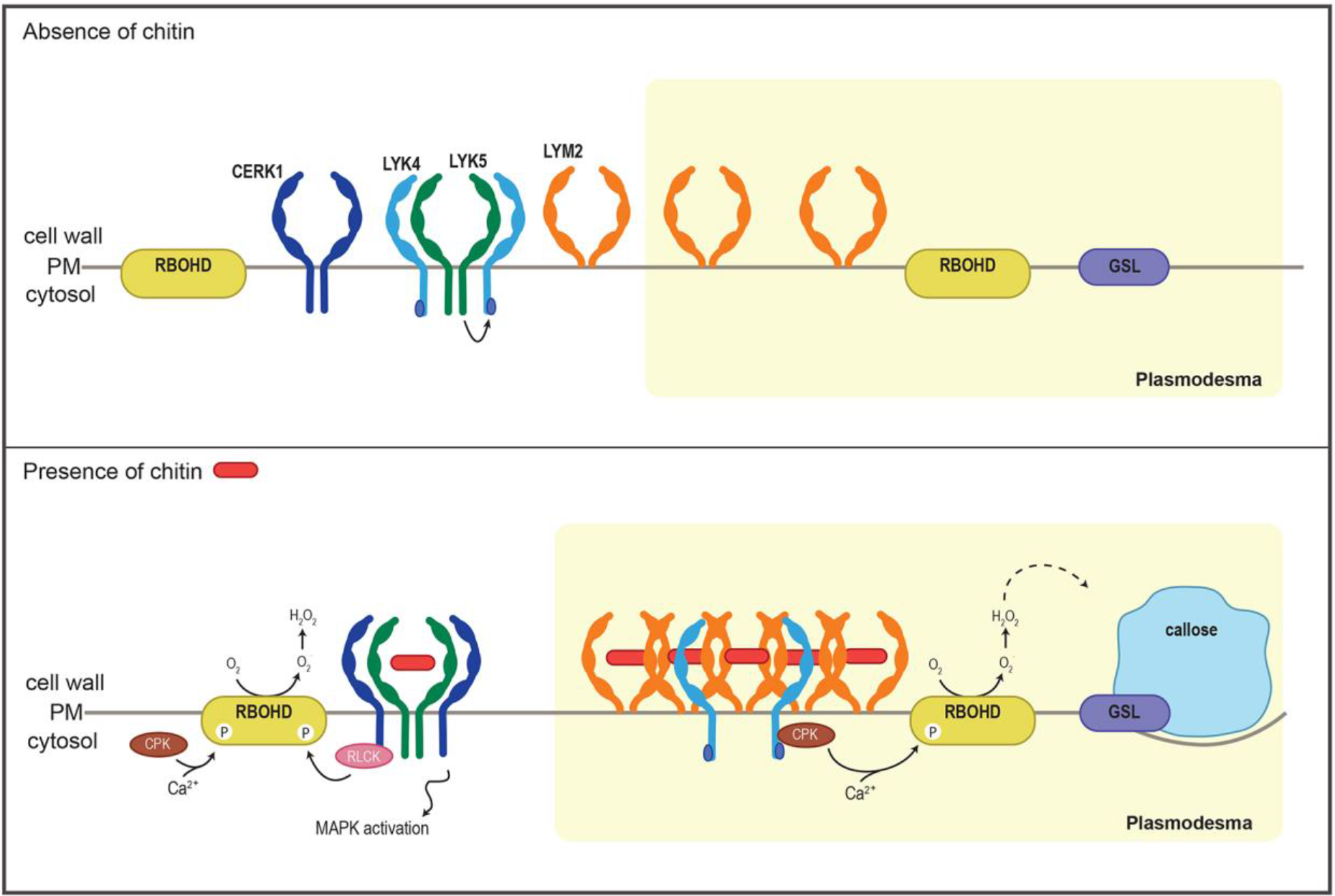
Model for LYM2-mediated chitin signalling in the plasmodesmal PM. This cartoon describes our proposal for some of the key elements of LysM signalling in the PM and plasmodesmal PM in response to chitin perception. The top panel represents functional, interactions under control conditions. Here, LYK5 (green) interacts with LYK4 (light blue) and mediates modification of LYK4 in the PM, and LYM2 (orange) is distributed throughout the PM and the plasmodesmal PM. In response to chitin (lower panel), LYK4 and LYK5 dissociate. LYK5 interacts with CERK1 (dark blue) to mediate signalling at the PM, and LYM2 accumulates at plasmodesmata where it forms a higher order complex. This complex recruits LYK4 and CPK6 (brown) which phosphorylates RBOHD (yellow) and induces callose (blue) synthesis via a glucan-synthase like enzyme (GSL, purple) to close PD.

This model is distinct from that for chitin perception in the PM in which a CERK1-LYK5 complex forms in response to chitin and triggers BIK1-dependent phosphorylation of RBOHD. Both the plasmodesmal PM and PM chitin-triggered signalling cascades activate RBOHD, but our analyses identified that CERK1 and LYM2 signalling function independently (Faulkner *et al.*, 2013). It is significant that while both chitin responses activate RBOHD there is no evidence of cross-talk between the ROS produced by CERK1 and LYM2. This independence suggests either that ROS signalling is highly localised. Alternatively, it is possible that the apoplastic environment surrounding the plasmodesmata insulates the plasmodesmal PM from external signals, but the observation that exogenously applied hydrogen peroxide can induce plasmodesmal responses (Fig. S5; Cui and Lee, 2016) suggests this is unlikely to fully explain the signalling localisation.

We have demonstrated that the plasmodesmal PM integrates signals independently of the PM and that plant cells are thus capable of exploiting receptor and signalling specificity to mediate localised and specialised responses. In the context of immune responses, this identifies that cells regulate cell-to-cell connectivity independently of other immune outputs. This work establishes that cell-to-cell connectivity and molecular exchange is a key component of how plant tissues respond to pathogen signals. Given that plasmodesmata are regulated in response to a range of abiotic and developmental signals it seems likely that the ability of a cell to independently regulate its plasmodesmata occurs in a variety of contexts and is a fundamental component of how tissues and organs respond as multicellular entities.

## Material and methods

### Plant materials

*Arabidopsis* plants were grown on soil in short days conditions with 10h light at 22 °C. *N. benthamiana* plants were grown on soil with 16h light at 23 °C. Arabidopsis Columbia-0 (Col-0) was used as wild-type plants. The mutant lines used in this study are: *rbohd* (Torres, Dangl and Jones, 2002), *lym2-1* (SAIL_343_B03, (Faulkner *et al.*, 2013)), *cerk1-2-*(GABI_096F09, (Miya *et al.*, 2007)), *lyk3* (SALK_140374 (Wan *et al.*, 2012)), *lyk4* (WiscDsLox297300_01C (Wan *et al.*, 2012)), *lyk5-2* (SALK_131911.31.20.x (Cao *et al.*, 2014)), *bik1* (Salk_005291C (Veronese *et al.*, 2006)), *cpk6-1* (SALK_093308) and *cpk6-2* (SALK_033392) (Mori *et al.*, 2006). RBOHD variants RBOHD_S339A/S343A_ and RBOHD_S39A/S339A/S343A_ are mutant variants of RBOHD transformed in to the *rbohd* mutant background as described (Kadota *et al.*, 2014). The RBOHD variants RBOHD_S133A_ and RBOHD_S163A_ were generated by complementation of the Arabidopsis *rbohd* mutant with constructs harbouring site specific mutations.

### DNA Constructs

Citrine-LYM2 is as previously described (Faulkner *et al.*, 2013). LYK4 and LYK5 coding sequences were PCR amplified from Col-0 cDNA and cloned into pDONR by Gateway BP reactions (Invitrogen). Entry Clones carrying LYK4 and LYK5 were verified by sequencing. To generate Cauliflower Mosaic Virus (CaMV) 35S promoter-driven LYK4 and LYK5 constructs C-terminally-tagged with GFP, RFP, HA or myc, pENTR clones were recombined by the Gateway LR reaction with pB7FWG2.0, pB7RWG2.0, pGWB14 and pGWB17 respectively. CPK6 was PCR cloned from Col-0 cDNA using gene-specific primers with Golden Gate compatible extensions (BpiI recognition sequences and fusion sites) and cloned into the universal acceptor plasmid pUAP1 by digestion/ligation using BpI1. Clones were validated by sequencing. Level 1 binary constructs were assembled in the pICH47751 vector with the following modules by digestion/ligation using BsaI: pUAP1 carrying CPK6; pICH51277 carrying CaMV 35S, pICSL50008 carrying GFP for C-terminal fusion, pICH41414 carrying CaMV 35S terminator. The BRI1-RFP fusion is as described (Bucherl *et al.*, 2017). For generation of RBOHD_S133A_ and RBOHD_S163A_ we used site-directed mutagenesis: DNA fragments were amplified by PCR using primers harbouring mutation from pBIN19g.pRbohD:3xFLAG-gRBOHD (Kadota *et al.*, 2014) and the mutated fragments were cloned between EcoRI and BamHI sites of pBin19g vector by using In-fusion enzyme.

### Transient expression in *Nicotiana benthamiana*

*Agrobacterium tumefaciens* GV3101 carrying the desired construct was cultured overnight and resuspended in MMA buffer containing 0.01M 2-(N-morpholino)ethanesulfonic acid (MES) pH 5.6, 0.01M MgCl_2_, 0.01M acetosyringone. Each *Agrobacterium* strain carrying the desired construct was mixed with an *Agrobacterium* strain carrying the P19 silencing suppressor. Each strain was adjusted to a final optical density of 0.2 at 600 nm and syringe-infiltrated into leaves of 4-week-old *Nicotiana benthamiana* plants. Material for experiments was harvested two days post-infiltration.

### Semi-quantitative RT-PCR analysis

RNA was extracted from leaves of 5-week-old Arabidopsis plants using RNeasy Mini extraction kit (Qiagen) and treated with Turbo DNA-free kit (Ambion) before cDNA synthesis using Reverse Transcriptase (Promega) according to the manufacturer’s instructions. Semi-quantitative RT-PCR was performed on the cDNA samples with primers listed in Table S2.

### Microprojectile bombardment assays

Microprojectile bombardment assays were performed as described (Faulkner *et al.*, 2013). 5- to 6-week-old expanded leaves of *Arabidopsis* plants were bombarded with 1nm gold particles (BioRad) coated with pB7WG2.0-GFP and pB7WG2.0-RFP_ER_, using a Biolostic PDS-1000/He particle delivery system (BioRad). The gold particles were prepared by precipitating 5 μg of each DNA plasmid construct on to the gold particles with 625mM CaCl_2_ (Sigma) and 10mM spermidine (Sigma). Bombarded leaves were infiltrated with 500 μg/mL chitin (mixture of chitin oligosaccharides, NaCoSy) or water 2 hours post bombardment. Bombardment sites were assessed 16 hours after bombardment by confocal (Leica SP5 or SP8) or epifluorescence microscopy (Leica DM6000) with a 25× water dipping objective (HCX IRAPO 25.0× 0.95 water). The number of cells showing GFP was counted for each bombardment site (marked by RFP_ER_). For each treatment, data were collected from at least 3 independent experiments, each of which consisted of leaves from at least three individual plants. The normality of each data set was assessed by a Shapiro-Wilk test. When normally distributed a Student’s t-test was applied and when the data was not normally distributed a non-parametric Mann-Whitney U test was used. Statistical significance was concluded when p-value was less than 0.05. Sample-specific details are listed in the figure legends.

### Protoplast preparation and transfection

Protoplasts were extracted and transfected as described previously (Yoo, Cho and Sheen, 2007). True leaves from 4-5-week-old Arabidopsis plants were cut in 1mm leaf strips using a razor blade. Leaf strips were immediately submerged into an enzyme solution containing 20mM MES (Sigma) pH 5.7, 1.5% (w/v) cellulase R10 (Yakult Pharmaceuticals, Japan), 0.4% (w/v) macerozyme R10 (Yakult Pharmaceuticals, Japan), 0.4M mannitol, 20mM KCl, 10mM CaCl_2_ and 0.1% bovine serum albumin (BSA). The enzyme solution was warmed to at 55 oC for 10min and filtered with 0.45μm syringe filter before use. Leaf strips were vacuum-infiltrated for 30 minutes in the dark and left at room temperature in the dark for 3 hours. The protoplasts in solution were then diluted with an equal volume of W5 solution containing 2mM MES pH 5.7, 154mM NaCl, 125mM CaCl_2_, 5mM KCl and filtered with a 75μm nylon mesh. Protoplasts were pelleted in a 30ml round-bottom Oak Ridge centrifuge tube at 150*g*, for 1min and re-suspended at 2×10^5^ml^−1^in W5 solution (the number of protoplasts was counted using a haemocytometer). Protoplasts were kept on ice for 30 minutes, pelleted at 150*g* and re-suspended at 2×10^5^ml^−1^in MMG solution containing 4mM MES pH 5.7, 0.4M mannitol and 15mM MgCl_2_, at room temperature before DNA/PEG/Calcium transfection. For transfection, 10μg of plasmid DNA was mixed gently with 100μl of protoplasts and 110μl PEG/Calcium solution containing 20% (w/v) PEG4000, 0.2M mannitol and 100mM CaCl_2_. The transfection mixture was incubated at room temperature for 5 minutes, diluted with W5 solution and centrifuged at 100*g* for 2min. Protoplasts were then re-suspended in WI solution containing 4mM MES pH 5.7, 0.5M mannitol and 20mM KCl and incubated at room temperature for about 16 hours. Transfected protoplasts in WI were centrifuged at 100g for 2min, the supernatant was removed, and the pellet was frozen on dry ice before being stored at −70 °C for further analyses.

### Microscopy

Confocal microscopy was performed on a Leica SP5, Leica SP8, or ZEISS LSM800 with a 25× water-dipping lens (HCX IRAPO 25.0 × 0.95 water), a 40× oil immersion lens (HCX PLAPO CS 40.0× 1.25 OIL), a 63× oil immersion lens (Plan-APOCHROMAT 63×/1.4 oil) or a 63×/1.2 water immersion objective lens (C-APOCHROMAT 63×/1.2 water). Citrine was excited with a 488 nm or 514 nm argon laser and collected at 525–560 nm. mRFP and mCherry were excited with a 561 nm DPSS laser and collected at 600-640 nm, GFP was excited with a 488 nm argon laser and collected at 505-530 nm, aniline blue was excited with a 405 nm UV laser and collected at 430-550 nm.

### Plasmodesmal callose staining and quantification

Plasmodesmal callose staining and quantification was performed as described (Xu *et al.*, 2017). The 8^th^ leaf of 5-6-week-old Arabidopsis plants was infiltrated with water or chitin. One hour after treatment, the leaf tissue was infiltrated with 0.01% aniline blue in PBS buffer (pH 7.4). Three areas were cut from one leaf and three different sites were imaged from each area. Callose deposits were imaged from the abaxial side of the leaf using a 63× oil immersion lens (Plan-APOCHROMAT 63×/1.4 oil) with a Leica SP5 confocal microscope. Aniline blue was excited with a 405 nm UV laser and collected at 430-550 nm. This was replicated for 5-12 leaves per genotype and treatment. Aniline blue stained plasmodesmal callose was quantified using automated image analysis pipeline “find plasmodesmata” written in Python (Xu *et al.*, 2017) (https://github.com/JIC-CSB/find-plasmodesmata). Three-dimensional segmentation was carried out by initially thresholding the input image (experimentally adjusted: default min 8000, max 15000). The resulting binary image was then segmented by connected component analysis. Post segmentation filtering was based on the number of voxels in each connected component. Objects <2 voxels (noise) and objects >100 voxels (callose accumulated in stomata) were removed. All annotated images were sanity checked prior to inclusion of data in the final set. The normality of each data set was assessed by a Shapiro-Wilk test. When normally distributed a Student’s t-test was applied and when the data was not normally distributed a non-parametric Mann-Whitney U test was used. Statistical significance was concluded when p-value was less than 0.05. Sample-specific details are listed in the figure legends.

### PD index (plasmodesmata/PM fluorescence intensity ratio)

Leaves of *N. benthamiana* transiently expressing the constructs of interest were observed 30 min after infiltration with chitin (500 μg/mL) or water (mock conditions) for PD index determination. The abaxial side of the leaf samples was imaged using a 63×/1.2 water immersion objective lens (C-APOCHROMAT 63×/1.2 water) with either a Leica SP8 or Zeiss LSM800 confocal microscope. PD index was determined by measuring the intensity values of Citrine-LYM2 at plasmodesmata and in two surrounding PM regions with ImageJ. PD index values were averaged for each image and 31 images for mock and chitin treatments were analysed by a Student’s t-test.

### Fluorescence Anisotropy

Leaves of *N. benthamiana* transiently expressing Citrine-LYM2 and cytosolic GFP were imaged with a Leica SP8 confocal microscope. Citrine was excited with a pulsed white light laser (488 nm) and emitted light was separated in to and perpendicular polarizations and detected by external SPADs with 500-550 nm filters. A series of 20 frames were merged and analysed using PicoQuant SymPhoTime 64. Anisoptropy (*r*) was calculated by

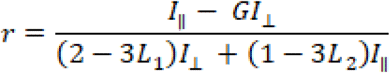

where G=0.481, L_1_=0.013 and L_2_=0.037. ROIs were defined that correlated to plasmodesmata and the PM for Citrine-LYM2 images or the cytosol for GFP images.

### FRET-FLIM analysis

Leaves of *N. benthamiana* transiently expressing the constructs of interest were used 30 min after infiltration with chitin (500 μg/mL) or water (mock conditions) for FRET-FLIM. The abaxial side of the leaf samples was imaged using a 63×/1.2 water immersion objective lens (Leica C-APOCHROMAT 63×/1.2 water). FLIM experiments were performed using a Leica TCS SP8X confocal microscope equipped with TCSPC (time correlated single photon counting) electronics (PicoHarp 300), photon sensitive detectors (HyD SMD detector), a pulsed laser (white light laser, WLL) with a range from 470 to 670nm. The WLL at 488 nm was used to excite GFP at 488nm. Laser power was kept low (0.5-3%) to avoid any bleaching of the samples. GFP emission was collected between 509-530nm. The repetition rate was set up to 40Mhz. Additional 488nm and 561nm notch filters were used to reduce interference reflected light. The instrument response function (IRF) was measured using erythrosine B as described (Stahl *et al.*, 2013). The FLIM data sets were recorded using the Leica LASX FLIM wizard linked to the PicoQuant SymPhoTime 64 software. The FLIM data sets were acquired by scanning each image until a suitable number of photon counts per pixel (minimum 1000) was reached. For the acquisition, the image size was set to 250 × 50 pixels with a speed of 50Hz, allowing a pixel dwell time of 19μs. The laser power was adjusted to reach a maximum count of 2000 kcounts per second. Each acquisition was stopped after 40 frames. Data were analysed by obtaining excited-state lifetime values of a region of interest (plasma membrane or plasmodesmata). Calculations were performed using the PicoQuant SymPhoTime 64 software instructions for FLIM-FRET-Calculation for Multi-Exponential Donors, using a two-exponential decay for GFP. The lifetimes were initially estimated by fitting the data using the Monte Carlo method and then by fitting the data using the Maximum Likely Hood Estimation (MLE). The amplitude weighted average donor lifetime with model parameter n=2 was used to calculate the average FRET-efficiency. FRET efficiency (E) was calculated by comparing the lifetime of the donor in the presence τDA or absence τD of the acceptor according to the following formula: 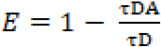. Data was analysed either by Student’s t-test for pairwise comparisons, or by ANOVA with Tukey-kramer post-hoc analysis for multiple comparisons. Statistical significance was concluded when p-value was less than 0.05. Sample-specific details are listed in the figure legends.

### FRAP analysis

For FRAP experiments we used leaves of *N. benthamiana* transiently expressing Citrine-LYM2, LYK4-RFP or LYK5-RFP treated for 30 min with chitin (500 μg/mL) or water (mock). FRAP was performed using a Leica TCS SP8X CLSM with a 63x/1.20 water immersion objective (Leica HC PL APO CS2 63×/1.20). To reduce scanning time the acquisition window was limited to 512 by 64 pixels. Citrine was excited at 514 nm and emissions detected between 527-550 nm. RFP was excited at 561 nm and emissions detected between 567-617 nm. ROIs were defined for plasmodesmata-localised Citrine-LYM2 and for PM-localised Citrine-LYM2, LYK4-RFP and LYK5-RFP. The FRAP protocol was as follows: 30 iterations were imaged at 0.095 sec/frame (pre-bleach); 15 (Citrine) or 60 (RFP) iterations at 0.095 sec/frame (bleach); and 50 iterations at 0.095 sec/frame followed by 120 iterations every 0.5 s (post-bleach). For bleach iterations the laser power was set to 100% and the FRAP booster was used.

Intensity data were normalised to the mean intensity of the first five frames and corrected for bleaching induced by pre- and post-bleach iterations. For the latter we collected non-bleached image acquisition decay curves. These data were themselves normalised to mean intensity of the first five frames and were fitted by a LOESS regression. Thus, all experimental FRAP data was corrected as follows:

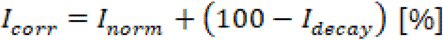

*I*_*corr*_ is the corrected intensity, *I*_*norm*_ is the normalised intensity, *I*_*decay*_ is the modelled intensity from the acquisition decay curve.

Data was further transformed to set the bleach base-line to zero: thus

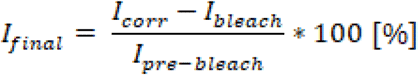

*I*_*final*_ is the final intensity, *I*_*bleach*_ is the intensity at the last bleach interation, *I*_*pre-bleach*_ is the mean of the pre-bleach values.

A LOESS curve was modelled to each dataset and each curve fit was sanity checked (Spira *et al.*, 2012). The intensity value at 60 s post-bleach was used as an approximation of the relative mobile fraction, adapted from Martiniere *et al.* (2012). Pairwise comparisons of estimated marginal means were performed in R 3.5.1 (R Core Team, 2018) using emmeans (Lenth et al., 2018; https://cran.r-project.org/web/packages/emmeans/index.html).

### Co-immunoprecipitation

Proteins were isolated using an extraction buffer containing 50 mM Tris-HCl pH 7.5, 150 mM NaCl, 5mM dithiothreitol (DTT), protease inhibitor cocktail (Sigma) 1:100, phosphatase inhibitor (Sigma) 1:200, 1 mM phenylmethanesulfonyl fluoride (PMSF) and 0.5% IGEPAL^®^ CA-630 (Sigma). For co-immunoprecipitation, GFP-Trap agarose beads (ChromoTek) were incubated with the protein samples for 2 hours at 4 °C with gentle agitation. Beads were then washed three times with a washing buffer containing 50 mM Tris-HCl pH 7.5, 150 mM NaCl, 5mM dithiothreitol, protease inhibitor cocktail 1:100, phosphatase inhibitor 1:200, 1 mM phenylmethanesulfonyl fluoride and 0.1% IGEPAL. After addition of Laemmli buffer (2×) containing 50 mM Tris-HCl pH 7, 5% SDS, 20% glycerol, 2% bromophenol blue, 5% β-mercaptoethanol, samples were boiled for 15 minutes at 95 oC. Protein samples were analysed by SDS-PAGE and transferred to Immuno-blot® PVDF membrane. Proteins were detected with α-GFP-HRP (Miltenyi Biotec, 130-091-833), α-Myc-HRP (Abcam, ab622928), α-HA-HRP (Abcam, ab173826), α-RFP-biotin (Abcam, ab34771) antibodies. α-RFP-biotin was detected using α-rabbit IgG–Alkaline Phosphatase (Sigma, a3687).

### Plasmodesmata extraction

The plasmodesmal purification method from Arabidopsis suspension cultures cells was modified for use with mature leaf tissue. Four expanded 5-week-old *N. benthamiana* leaves, transiently expressing the desired construct, were ground in liquid nitrogen to a fine powder. The powder was ultrasonicated in extraction buffer (50 mM Tris-HCl pH 7.5, 150 mM NaCl, 1× cOmplete™ ULTRA EDTA-free protease inhibitors (Roche), 1 mM PMSF, 1% w/v PVP-40kDa (Sigma)), followed by high-pressure homogenisation (EmulsiFlex B15, Avestin) to produce the “Total” fraction. Triton-X100 (0.5% v/v) was added to the resultant homogenate and the sample was centrifuged at 400*g* at 4°C to collect a crude cell wall extraction. Cell walls were washed five times with extraction buffer, and once in cellulase buffer (20 mM MES-KOH pH 5.5, 100mM NaCl, 1× cOmplete™ ULTRA EDTA-free protease inhibitors (Roche), 1 mM PMSF, 4.4% w/v mannitol). Washed cell wall material was resuspended in an equal volume of cellulase buffer with 2% (w/v) Cellulase R-10 (Yakult pharmaceuticals, Japan) and shaken at 37°C for one hour. Undigested cell wall material was removed by centrifugation at 2500*g* at 4°C. The supernatant was ultracentrifuged at 130,000*g* at 4°C to collect plasmodesmal membranes. The membrane pellet was resuspended in 50 μL resuspension buffer (50 mM Tris-HCl pH 7.5, 150 mM NaCl, 5 mM DTT, 1× cOmplete™ ULTRA EDTA-free protease inhibitors (Roche), 1 mM PMSF, 0.2% v/v IPEGAL^®^ CA-630 (Sigma)) as a “PD” fraction. Protein extracts were boiled in Laemmli buffer and separated by 10% SDS-PAGE. Proteins were transferred to Immuno-blot^®^ PVDF (0.2 μm, Bio-Rad) by semi-dry transfer. The membrane was probed with primary-conjugated HRP antibodies against HA (1:5000, ab173826, Abcam) and Myc (1:5000, ab62928, Abcam). HRP signal was developed with SuperSignal West Femto (ThermoFisher) and detected using ImageQuant LAS 500 (GE Healthcare).

### ROS burst measurement

Leaf discs (4 mm in diameter) were immerged in in 96-well white plates (Greiner bio-one) containing sterile water overnight in the dark. The next day, the water was replaced by a solution containing 20 μg/mL HRP (Sigma), 17 μg/mL luminol (Sigma), and chitin (NaCoSy, 500 μg/mL) or water (mock). Integrated luminescence was measured using a luminometer (Varioskan Flash) for 40 minutes. The normality of each data set was assessed and confirmed by a Shapiro-Wilk test. Thus, a Student’s t-test was applied for pairwise comparisons and significant differences were concluded when p-value was less than 0.05. Sample-specific details are listed in the figure legends.

### Statistical analyses

Statistical analyses were performed using Genstat^®^ version 18 or R 3.5.1. Details are in the relevant methods section and/or figure legends. n-values are detailed in Table S1.

## Acknowledgements

The authors acknowledge access to the JIC Bioimaging Facility we thank the staff for their assistance and training with the microscopes. The authors particularly thank Grant Calder and Yvonne Stahl (University of Düsseldorf) for assistance in developing the FRET-FLIM methods. *lyk3*, *lyk4* and *lyk5-2* seeds were provided by Gary Stacey. This work was funded by: the Biotechnology and Biological Research Council (BB/L000466/1, CF; BBS/E/J/000PR9796, CF); the European Research Council (grant ‘INTERCELLAR’, CF); and the Gatsby Charitable Foundation (CZ). The authors thank Caroline Dean for discussions and critical reading of the manuscript.

## Supplementary Data

**Figure S1.**
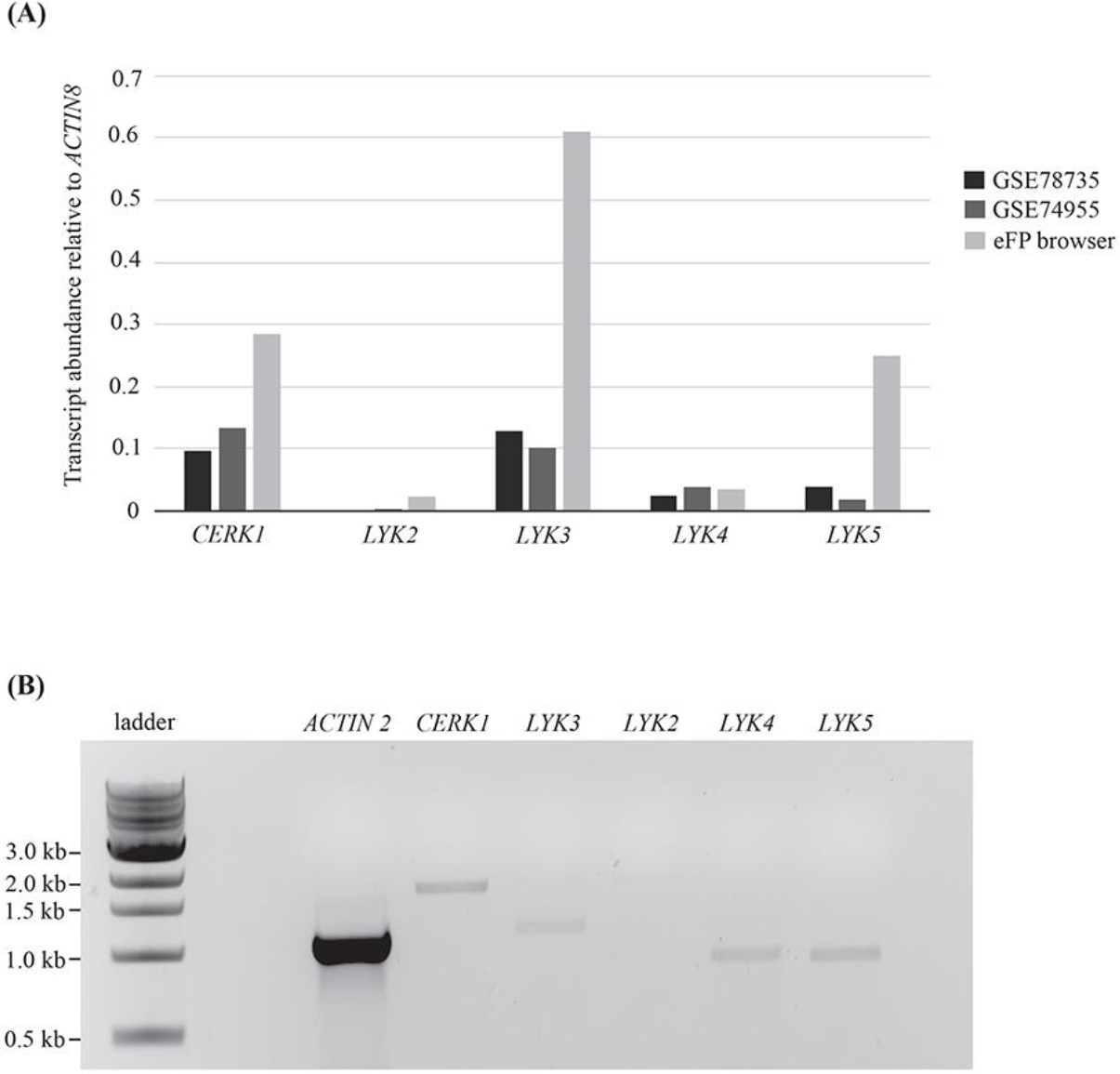
*LYK2* is not detected in mature leaves. (A) Publically available gene expression data from RNAseq (GSE78735 Hillmer *et al.,* 2017; GSE74955, Yamada *et al.,* 2016) and microarray experiments (eFP browser, Winter *et al.*, 2007) showing mRNA abundance of *LYK* family members relative to *ACTIN8*. The experimental data sets are indicated. (B) Semi-quantitative transcript abundance of *LYSM-RKs* in 5–6‐week‐old Arabidopsis leaf tissue. Transcript abundance was detected by RT-PCR and *ACTIN2* was used as an internal control. The primers used for RT-PCR are listed in the supplemental data (Table 2). RT-PCR reaction with these primers give the following sizes: ACTIN2, 1131bp; CERK1, 1854bp; LYK2, 1960bp; LYK3, 1275bp; LYK4, 1083bp; LYK5, 1082bp.

**Figure S2.**
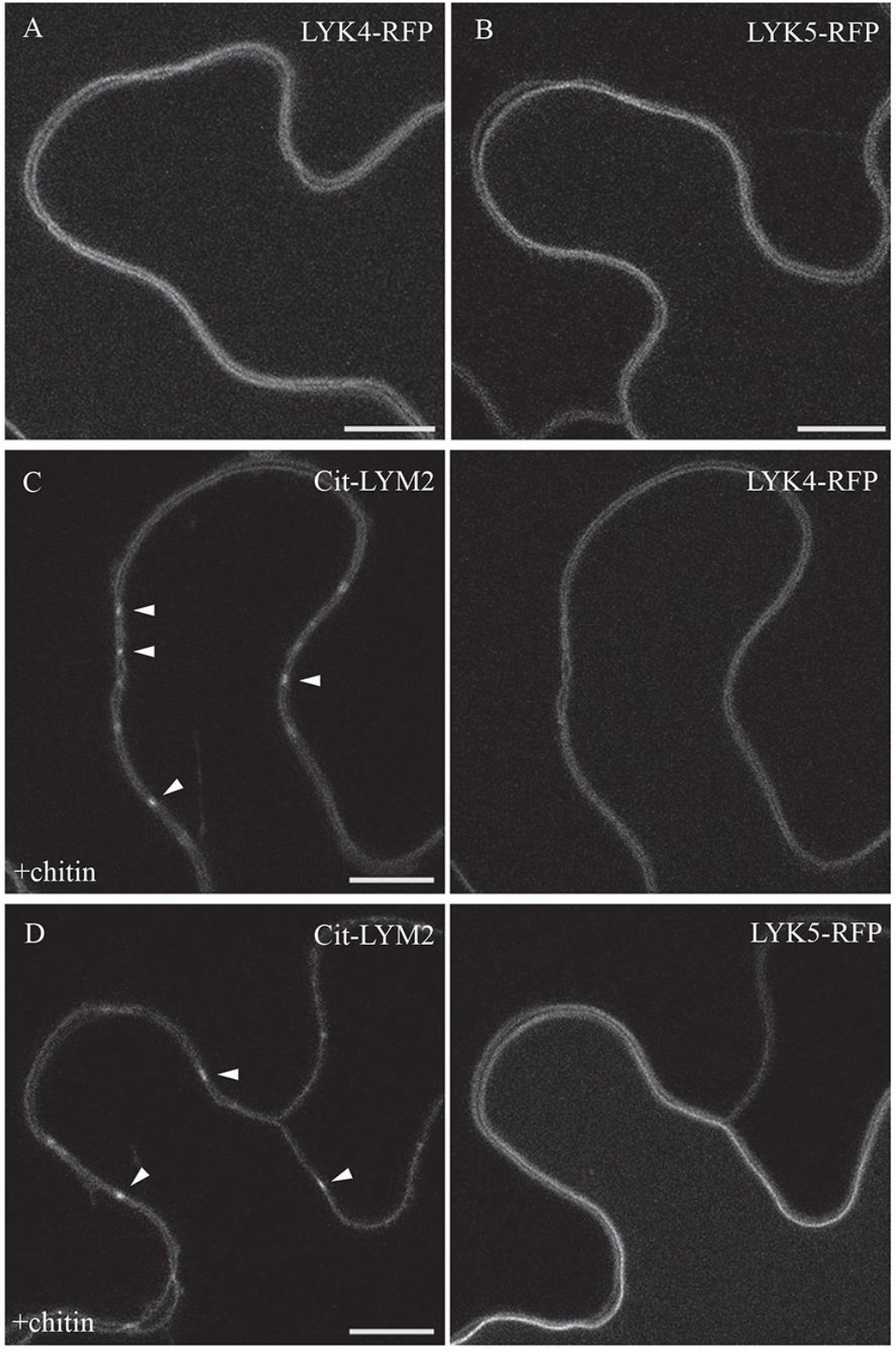
LYK4 and LYK5 localise to the PM. LYK4-RFP (A) and LYK5-RFP (B) localise evenly across plasma membrane when transiently expressed in *N.benthamiana*. (C) and (D) show localisation of LYK4-RFP and LYK5-RFP respectively in tissue that is co-expressing Citrine-LYM2 (Cit-LYM2) and has been treated with chitin for 30 min. Arrowheads identify Citrine-LYM2 marked plasmodesmata. Scale bars are 10 μm.

**Figure S3.**
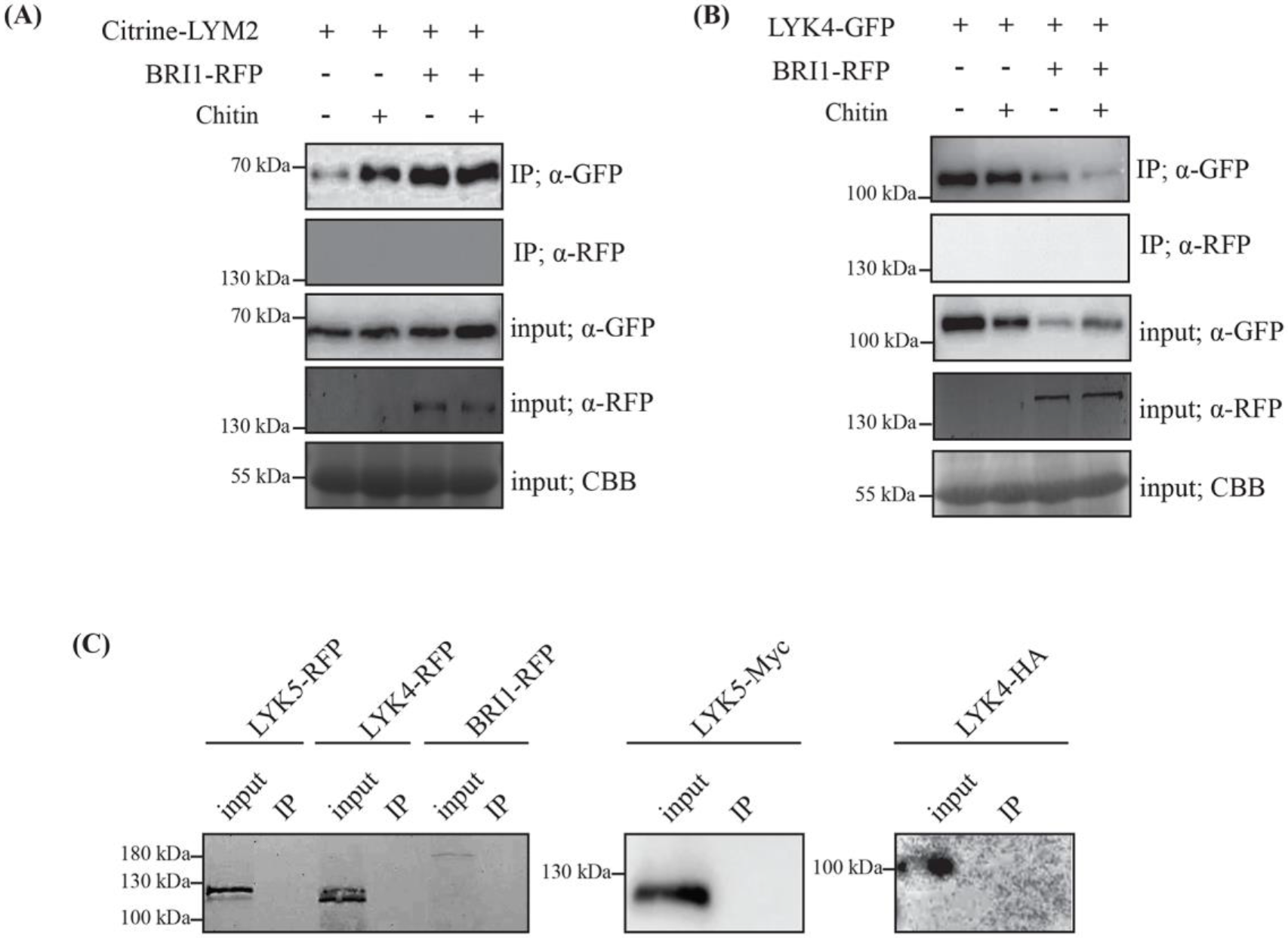
Negative controls for immunoprecipitation experiments. (A) and (B) Western blot analysis of immunoprecipitated protein extracts from *N. benthamiana* tissue expressing BRI1-RFP and either Citrine-LYM2 (A) or LYK4-GFP. (B) Citrine-LYM2 and LYK4-GFP were immunoprecipitated using GFP-trap beads and Western blots were probed with α-RFP to detect BRI1-RFP. Coomassie brilliant blue stained membranes serve as a loading control (CBB). Experiments were repeated three times with similar results. (C) LYK5-RFP, LYK4-RFP, BRI1-RFP, LYK5-Myc, LYK4-HA were immunoprecipitation with GFP-trap beads. Western blots of the input and immunoprecipitated (IP) samples were probed with antibodies α-RFP, α-Myc and α-HA.

**Figure S4.**
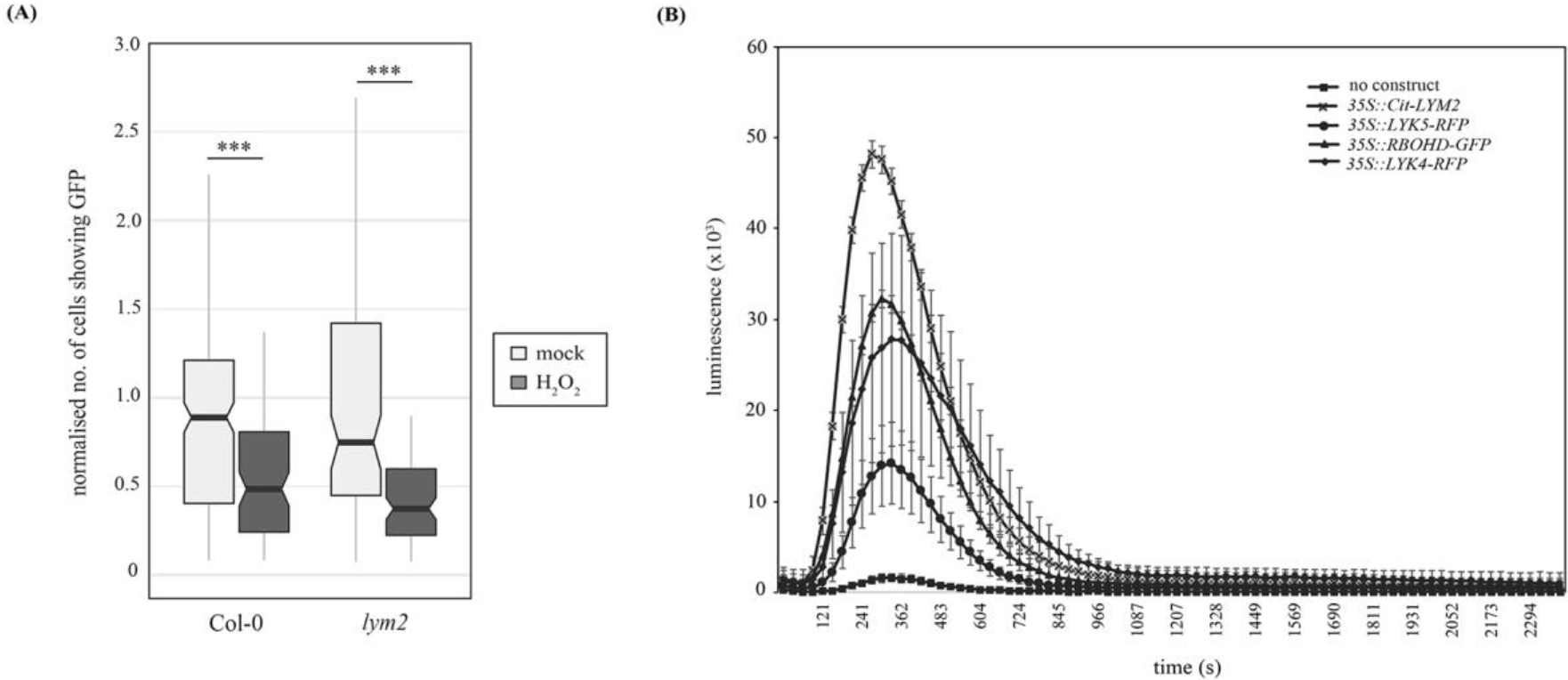
LYM2 acts upstream of ROS signalling in plasmodesmata closure. (A) Microprojectile bombardment into leaf tissue shows that H_2_O_2_ induces a reduction in GFP movement between cells in both Col-0 and *lym2-1* leaves. The number of cells showing GFP has been normalised to the mean of the mock data within genotypes. This data is summarised in box-plots in which the line within the box marks the median, the box signifies the upper and lower quartiles, the minimum and maximum within 1.5 × interquartile range. Data was analysed by a Students’ t-test, or by a Mann-Whitney U-test for non-normal data (n ≥ 93, ***p-value < 0.001). (B) Total accumulation of ROS in chitin-treated N. benthamiana leaf discs transiently expressing Citrine-LYM2, LYK4-RFP, LYK5-RFP, or RBOHD-GFP. Mean RLU are plotted over time (s). Error bars are SEM.

**Figure S5.**
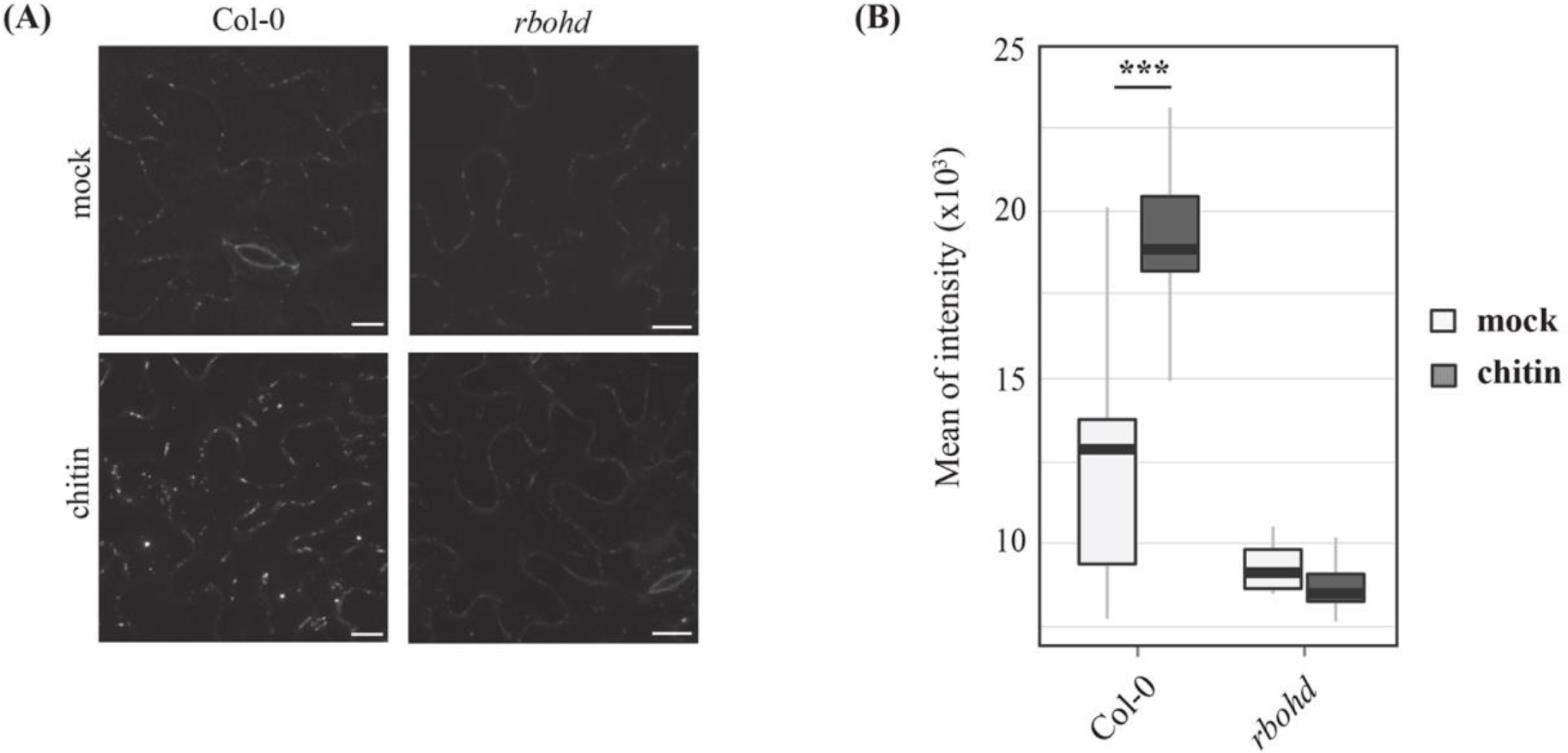
*rbohd* mutants do not deposit callose at plasmodesmata in response to chitin. (A) Confocal images of aniline blue stained plasmodesmal callose deposits in leaf tissue of 5–6‐week‐old leaves of Col-0 and *rbohd* mutant plants. Images were acquired 30 min post-infiltration with water or chitin. Scale bars are 15 μm. Images for Col-0 are as presented in Fig. 1B. (B) Quantification of plasmodesmata-associated fluorescence of aniline blue stained callose using automated image analysis. In contrast to Col-0, the *rbohd* mutant does not show increased aniline blue fluorescence at plasmodesmata in response to chitin. This correlates with the flux phenotype and identifies that chitin-triggered plasmodesmata closure is caused by callose deposition at plasmodesmata. The fluorescence intensity is summarised in box-plots in which the line within the box marks the median, the box signifies the upper and lower quartiles, the minimum and maximum within 1.5 × interquartile range. Data was analysed by pairwise Students’ t-test comparing mock to chitin treated tissue within genotypes (n ≥ 31, ***p-value < 0.001). Data for Col-0 is as presented in Fig. 1B.

**Table S1.**
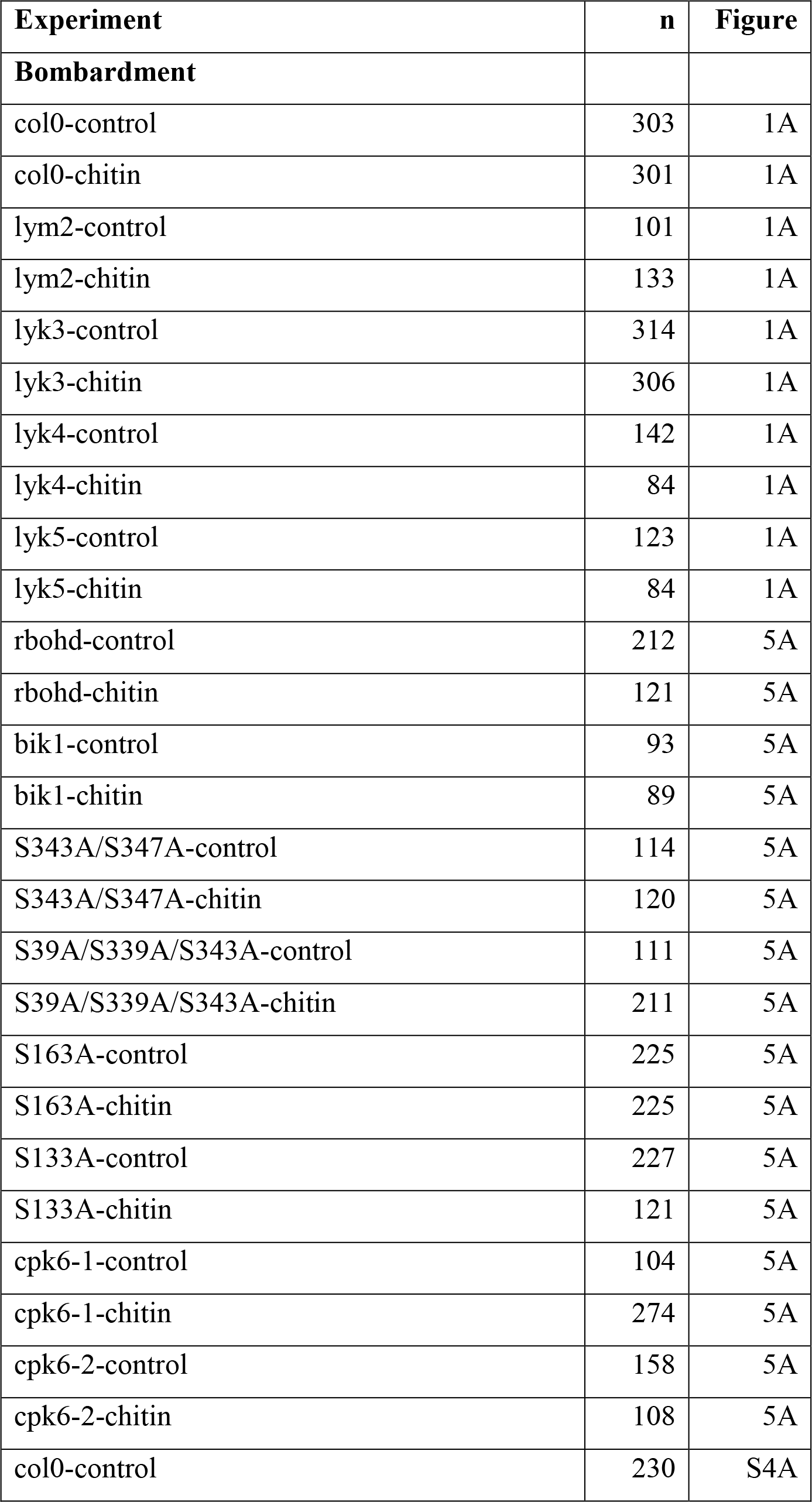
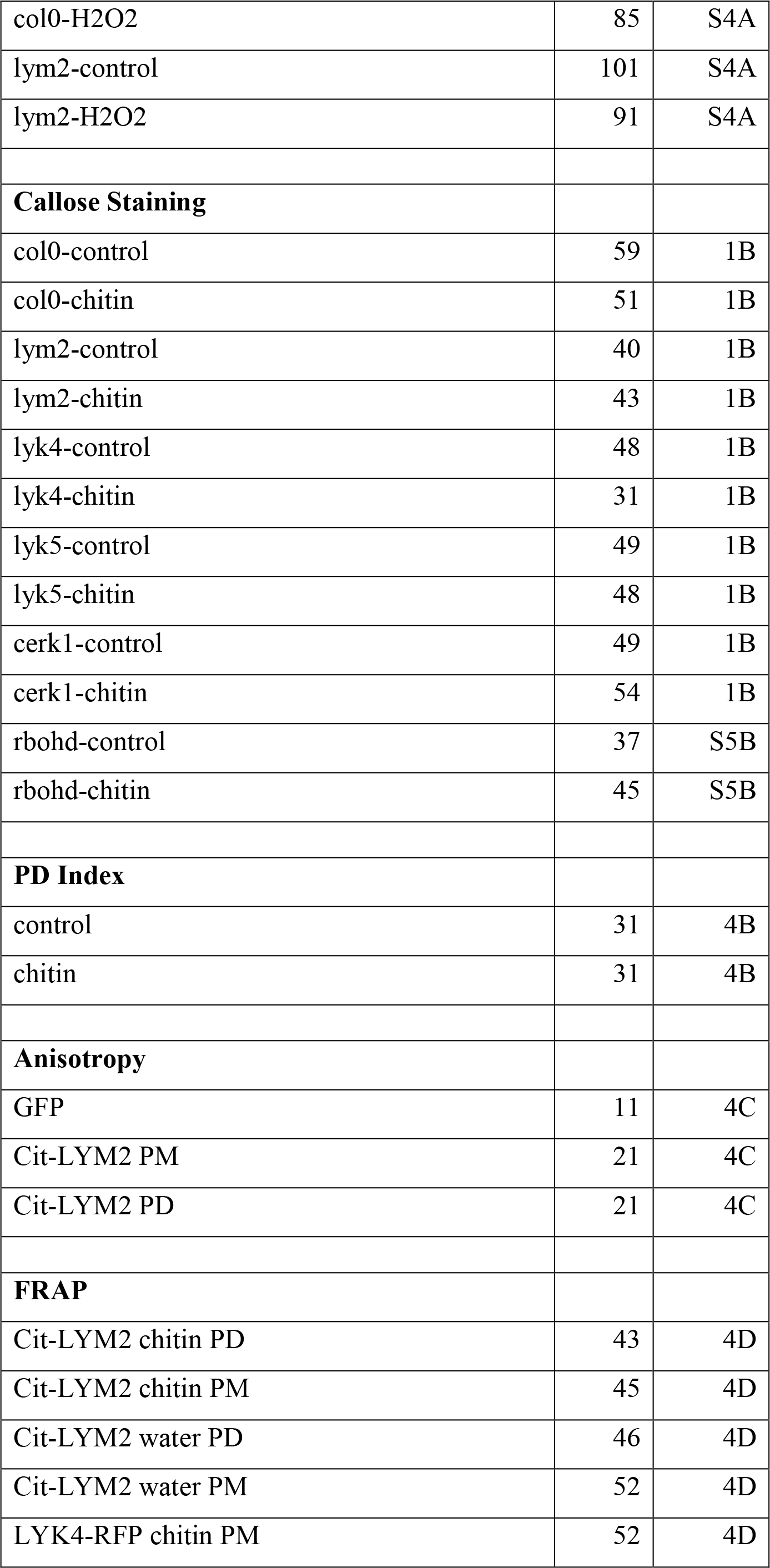
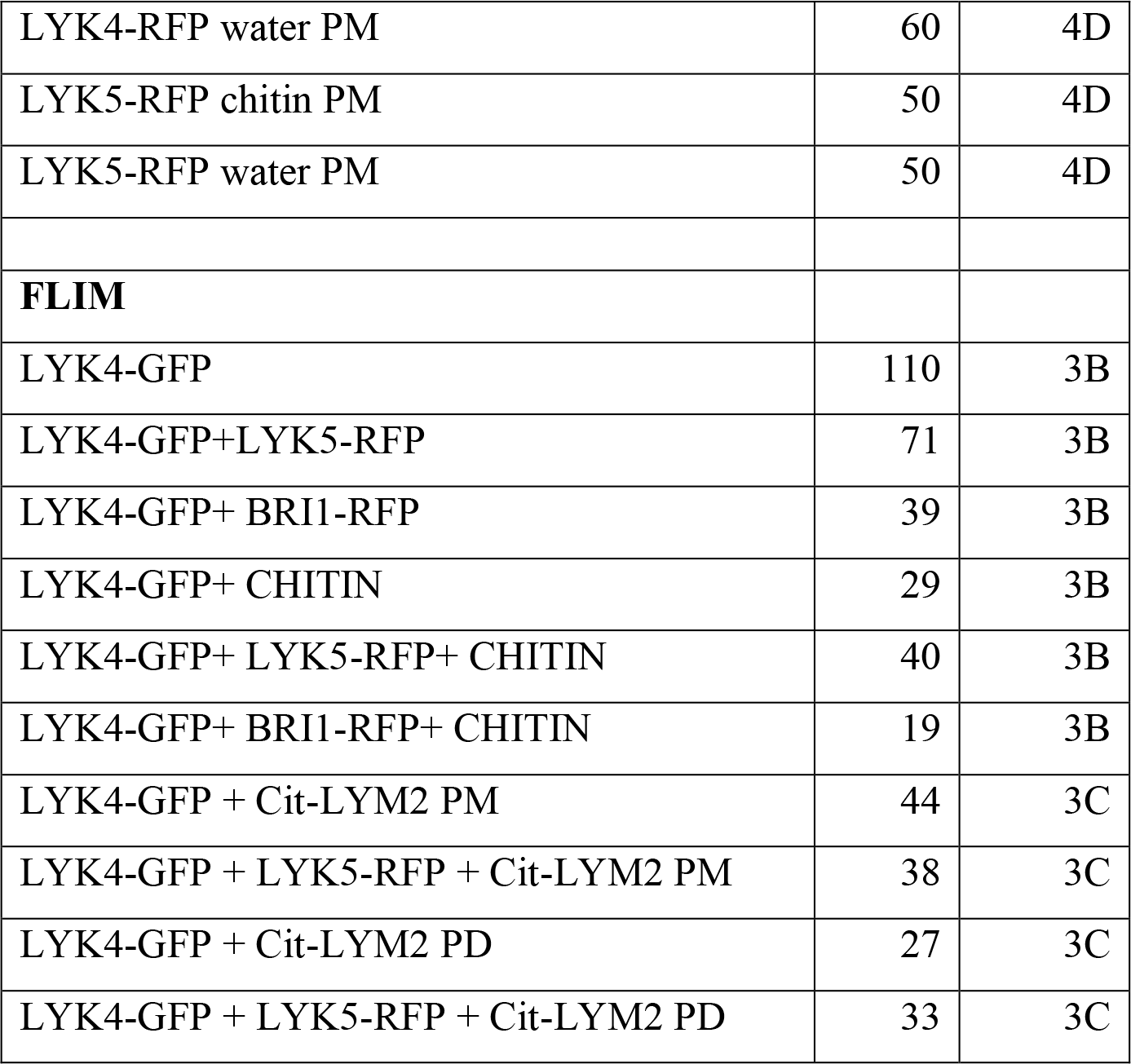
n-values for experiments in which different samples had different numbers of replicates.

